# Quantitative guiding of developmental cell fate transitions using a dynamical landscape model

**DOI:** 10.1101/2024.11.16.623539

**Authors:** Ismail Hajji, Francis Corson, Wolfgang Keil

## Abstract

During development, cells gradually assume specialized fates via changes of transcriptional dynamics in thousands of genes. Landscape modeling approaches, which abstract from the underlying gene regulatory networks and reason in a low-dimensional phenotypic space, have been remarkably successful in explaining terminal fate outcomes. However, their implications for the dynamics of fate acquisition have so far not been tested *in vivo*. Here we combine a landscape model for *C. elegans* vulval fate patterning with temporally controlled perturbations of EGF and Notch signaling in vivo using temperature-sensitive mutant alleles. We show that this approach quantitatively predicts non-intuitive fate outcomes and pathway epistasis effects. We further infer how cell fate transitions can be guided towards specific outcomes through timed pulses of signaling activity and verify these model predictions experimentally. Our results highlight the predictive power of landscape models and illustrate a new approach to quantitatively guide cell fate acquisition in a developmental context.

## Introduction

During development, cells proceed through a series of decisions between alternative cellular states to ultimately reach their final fates within the organism. Understanding, predicting, and manipulating cell fate has been a long-sought goal of developmental and regenerative biology. However, the determinants of cellular identities are exceedingly complex. The specification of different cell types from initially equivalent progenitors relies on the response to time-dependent signals, interpreted in a context-dependent manner through signaling cascades and gene regulatory networks that often involve dozens of molecular players. Recent technological advances are making it possible [1–5] to capture at single-cell resolution the genome-wide changes in transcriptional programs and epigenetic states that accompany cell state transitions, but deriving from such data an intuitive and predictive understanding of cell fate specification remains an open challenge.

C. H. Waddington contrasted the overwhelming complexity of molecular mechanisms with his nowadays omnipresent metaphor of the epigenetic landscape, in which developmental trajectories are likened to the motion of a ball rolling down a hill, with different valleys representing different fates. The landscape itself was envisaged as controlled by an underlying, web-like network of genes. In its essence, this metaphor captured the emergence of simplicity (a limited number of developmental outcomes) from a complex set of interacting molecular mechanisms. Giving a concrete mathematical form to this idea, several recent studies have shown that ‘geometric’ or landscape models - simple dynamical models that abstract away much of the underlying molecular complexity - allow both a general categorization of possible cellular decision structures, and an intuitive description and quantitative predictions for fate patterning in concrete systems [6–10]. In this approach, the general structure of the landscape, and a minimal parametrization, is postulated according to the known biology of a given system (e.g., so many valleys for so many possible fates), and the requisite quantitative parameters are then fit to experimental data. Although it ignores much of the underlying molecular details, this approach has been successfully applied to diverse systems, from pattern formation in a small group of cells in *Caenorhabditis elegans* (*C. elegans*) vulva [6,7], to self-organized patterning in *Drosophila* [8], metazoan segmentation [11], and the differentiation of mouse embryonic stem cells (mESCs) *in vitro* [12].

As a rather minimal instance that nonetheless embodies many general aspects of multicellular patterning, *C. elegans* vulval patterning provides an ideal context to pursue the implications of geometric models. The vulva is formed from a row of six vulval precursor cells (VPCs, called P3.p-P8.p) in the second and third larval stages (Fig. 1A). Each of the six VPCs has the potential to adopt any one of three different cell fates (called 1°, 2°, and 3°). In the wild type (WT), an inductive signal originating from the anchor cell (AC) of the somatic gonad induces three cells (P5-7.p) in an invariant 2°1°2° pattern. These cells undergo three rounds of divisions to generate the vulval tissues, while the remaining three cells adopt the uninduced 3° fate, and fuse with the epidermal syncytium after one division (the complete fate pattern can thus be encoded as 3°-3°-2°-1°-2°-3°, or 332123 in short) (Fig. 1B). In half of WT individuals, P3.p does not become receptive to inductive signals and fuses to the hypodermal syncytium prior to the induction time window. Vulval cell fates are specified via two conserved signaling pathways [13–18]: (i) the Ras/MAPK signaling pathway, activated by an Epidermal Growth Factor (EGF)-like ligand called LIN-3, secreted by the AC and (ii) lateral Delta-Notch signaling between neighboring VPCs via transmembrane (APX-1 and LAG-2) and diffusible (DSL-1) Notch ligands [19,20]. Briefly, EGF induces the 1° fate, which promotes expression of Notch ligands primarily in P6.p, inducing the 2° fate and repressing the 1° fate in the neighboring cells, P5.p and P7.p.

**Figure 1:**
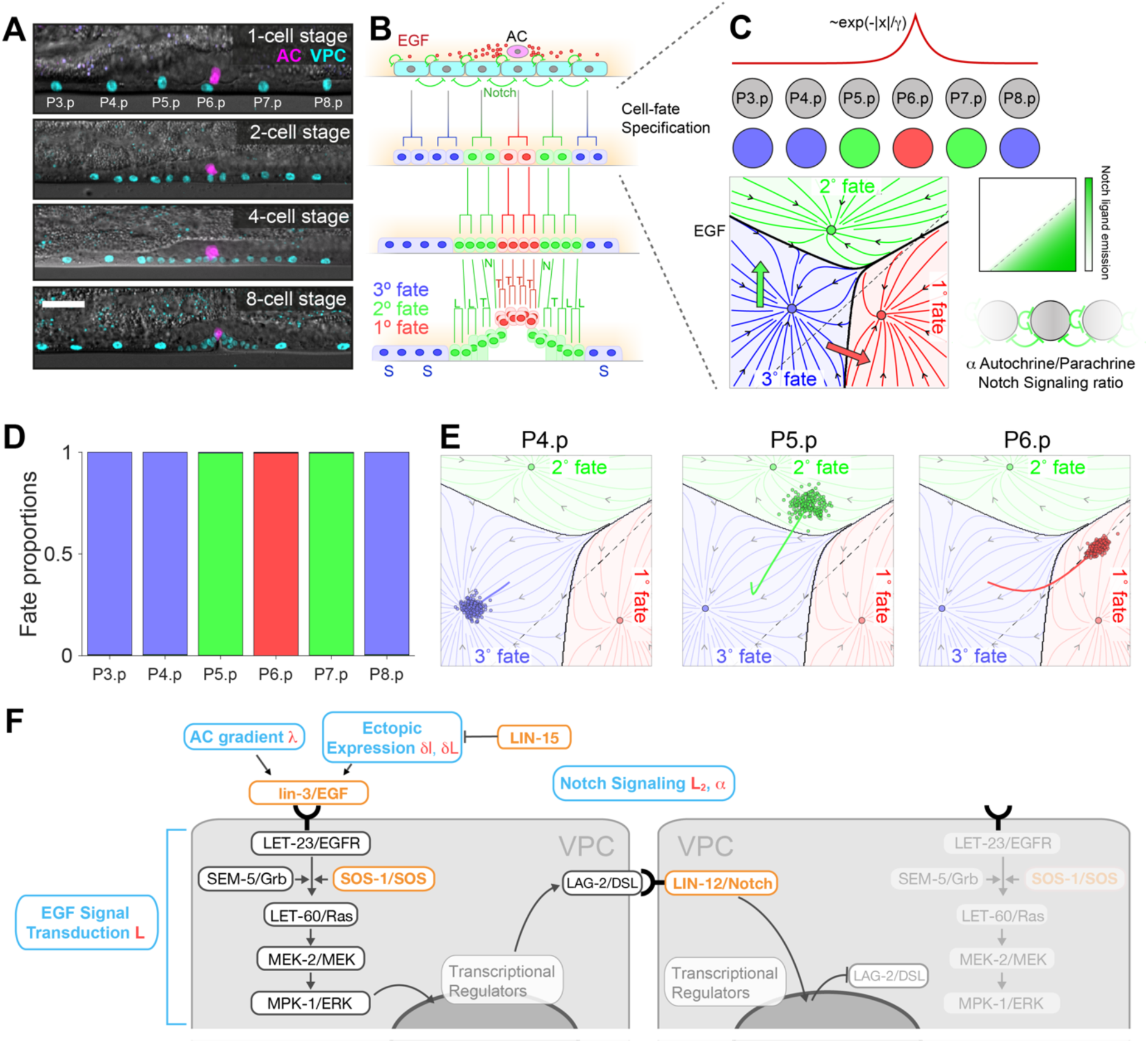
Cell fate acquisition during *C. elegans* vulval development. (**A**, **B**) Micrographs (**A**) and schematics (**B**) of *C. elegans* of vulval development. Scale bar, 10um. (**A**) The transgene *arTi85* [*lin-31p*::*ERK-KTR*(*NLS3*)*-mClover-T2A-mCherry-H2B*::*unc-54 3’UTR*] (1) (cyan) labels vulval precursor cells and descendants throughout all stages of vulval development, *qyIs362 [lin-29A/B>GFP]* (magenta) brightly labels the anchor cell starting from the mid L2 larval stage. Together, they allow accurate location of VPCs and scoring of vulval fate induction. (**B**) The vulva forms from a row of six precursor cells (VPCs), P3.p–P8.p, which are competent to adopt three possible fates, inner vulval or primary (1°, red), outer vulval or secondary (2°, green), and non-vulval or tertiary (3°, blue). Fates are identified by characteristic cell division patterns (the orientation of terminal divisions is indicated by letters: L, longitudinal; T, transverse; N, undivided; S, fused with the epidermal syncytium; underlined cells are adhered to the cuticle). In WT, a graded EGF signal from the anchor cell and lateral Notch signaling among VPCs (green arrows) specify an invariant pattern of fates, 32123. (**C**) Top, Our landscape model assumes an exponential EGF gradient originating from the AC, and autocrine and paracrine Notch signaling produced by the VPCs according to their current state. Bottom, Two-dimensional fate plane of the landscape model for vulval fate specification [6,7]. Black dots indicate valleys of the landscape, and black arrowheads the direction of cell state flow. Colored lines indicate cell fate basins (red, 1° fate; green, 2° fate; blue, 3° fate), separated by black lines. The dashed line is where Notch ligand expression is half maximum (see also green color code in inset, right). (**D**) Fate proportions for the six VPCs in the model for WT. (**E**) Model fate plan and trajectories for WT VPCs; lines are average trajectories of P4.p (blue), P5.p (green), and P6.p (red) over (200 realizations). Shaded regions show the flows at the end of the competence period into the three valleys for fates 1°-2°-3°. Dots show positions of individual model realizations after the competence period (colored according to the WT fate of each cell as per **C**), after which they follow the arrows into the valleys for fates 1°-2°-3°. (**F**) Diagram of the EGF and Notch signaling pathways. The colored boxes show where we intervene in the signaling pathways (orange) and the corresponding model parameters in Eqs. 2-3 (red).

A simple geometric model for vulval patterning, in which each cell travels in a two-dimensional landscape with three valleys (fates) was introduced in [6,7]. The landscape is independently tilted by EGF and Notch as development proceeds (Fig. 1C, Suppl. Movie 1, see Materials & Methods for details). The initially equivalent VPCs start from the same location at the beginning of competence, but different signaling histories take them along different trajectories. The model also incorporates a stochastic term to allow for the inherent variability of cellular dynamics. The states of the VPCs in multiple repeats of a simulation can be represented as clouds of points traveling in the plane (Fig. 1E). Whereas in the wild type, each of these clouds lies well within a valley at the end of competence, corresponding to an invariant fate pattern (Fig. 1D, Suppl. Movie 2), perturbations affecting signaling can cause one or several clouds to lie across a boundary between valleys, accounting for partially penetrant phenotypes (variable fate patterns). This model reproduces a wide variety of VPC fate patterns of animals defective in the two signaling pathways. Notably, phenotypic epistasis arising from geometric effects alone could explain several non-intuitive cell fate outcomes that had previously been attributed to context-dependent pathway action via downstream effector switching [21] or direct inhibitory interactions between the EGFR-MAPK and Notch pathways [19,22,23].

Since geometric models for developmental patterning explicitly model the *dynamics* of cellular fate trajectories (Fig. 1E, Suppl. Movie 1), they also make quantitative predictions for the impact of dynamical perturbations (e.g., of cellular signaling) *during* the cell fate acquisition process. While this is one of their most powerful features, such predictions remain to be tested *in vivo*. The *in vitro* experiments on mESCs performed in [12], while giving close to optimal external control over the signaling dynamics, hardly capture the full complexity of *in vivo* development, since secretion of the endogenous ligands was blocked. Thus, up to now, dynamical predictions of geometric models for multicellular patterning have not been challenged with quantitative *in vivo* measurements and perturbations *during* the differentiation process.

Here, we accomplish such tests, using vulval specification in *C. elegans* as a model system. We first construct combinations of mutants of the EGF/Ras/MAPK and Notch pathways with several alleles being strongly temperature sensitive. We show that the landscape model developed in [6,7], based on signaling parameters fit for the individual mutants, quantitatively captures these new allelic combinations at various growth temperatures. In doing so, we uncover new epistasis effects in the data that the model quantitatively explains without explicit pathway interactions. We then employ the temperature sensitivity of the allelic combinations to manipulate cell signaling during vulval cell fate acquisition using timed episodes of high and low cultivation temperatures. This allows us to ‘guide’ developmental trajectories towards predicted fate patterns, such as the maximal rescue of a mutant allele via a secondary mutation. The excellent agreement between the experimentally observed cell fate statistics and our theory demonstrates a general strategy to predictively control multicellular fate patterning *in vivo*.

## Results

### Landscape parametrizations of genetic perturbations to the EGF and Notch signaling pathways

Our goal is to quantitatively guide *in vivo* vulval cell fate phenotypes in predictive ways by combining temperature-sensitive genetic perturbations of EGF/MAPK and Notch signaling activity with geometric modelling of the underlying cell fate acquisition dynamics. Throughout this paper, we use the mathematical description of the cell-state landscape proposed in [6,7] to model *C. elegans* vulval fate acquisition. In that description, the current state of each cell is represented by a position vector 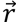 in a two-dimensional ‘fate plane’, whose time evolution is given by

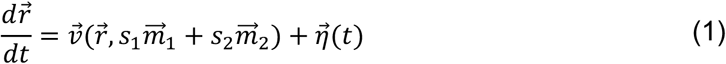

In this equation, the vector field 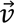 (detailed in Materials & Methods) describes the ‘landscape’ in which the cell is ‘moving’, and its dependence on the EGF/MAPK and Notch signaling activity levels, *s*_1_ and *s*_2_, which in WT we equate with the ligand levels received by the cell, *l*_1_ and *l*_2_. The scalars *s*_1,2_ multiply vectors 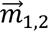 that are fixed for all genetic backgrounds. In addition to this deterministic term, the stochastic term 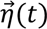 builds in the inherent variability of cell dynamics and accounts for the partially penetrant phenotypes (variable outcomes) observed in many conditions. A strong assumption of the model is that the effect of arbitrary signal levels is captured by the linear combination 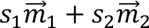, which can be thought of as the ‘force’ with which the two signals ‘push’ the cell, in directions controlled by the vectorial parameters 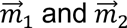 (see Fig. 1C, Suppl. Movie 1). Thus, combined perturbations affecting the EGF/MAPK and Notch pathways (as in double mutants) are described by just combining the corresponding changes in the model parameters that represent ligand levels and signal transduction efficiency (as depicted in Fig. 1F). Absent any explicit pathway coupling in the model, epistatic effects solely arise from the interplay between signaling responses and the underlying, nonlinear landscape. The morphogens act during a competence period after which cells relax into valleys corresponding to fates 1°-2°-3°.

For the EGF/MAPK pathway, WT ligand levels from the AC are described by a time-independent exponential gradient, i.e., a vector of ligand levels *l*_1_ = {*γ*^3^, *γ*^2^, *γ*, 1, *γ*, *γ*^2^} for the six cells, where the parameter *γ* < 1 controls the sharpness of the gradient (see Materials and Methods) [6,7]. The perturbations considered here include mutations that affect ligand production by the AC, mutations that results in ectopic ligand production, and mutations affecting signal Ras/MAPK transduction efficiency. For the most general perturbation, the signal levels received by the six cells are given by

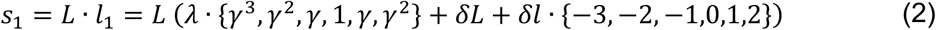

Here, the prefactor *L* represents signal transduction efficiency in the ERK pathway (e.g. SOS-1 in Fig. 1F) that modulates the response of all cells to all sources of ligands (*L* = 1 for WT and *L* < 1 for mutants in which signal transduction is impaired). Mutations affecting ligand production by the AC are represented by multiplying the WT gradient by a factor *λ*, whereas mutations that result in ectopic ligand production are represented by an additive term; experiments considered in the following require that we allow for non-uniform ectopic ligand levels, which for simplicity are assumed to exhibit a linear gradient, with offset *δL* and slope *δl*.

As regards lateral Notch signaling, on the other hand, the emission of Notch ligand by each cell is a function its current state, described by a scalar function 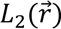. The Notch ligand level *l*_2_ to which each VPC is exposed includes contributions from its neighbors and from autocrine signaling. The only perturbation considered here is a mutation in *lin-12*/Notch that results in constitutive (ligand-independent) activation of the receptor, described as an additive contribution to signaling activity. Thus, in the most general case the Notch signaling activity of a cell (e.g. P6.p) is

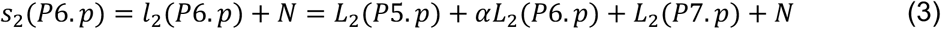

where *α* is the relative strength of autocrine vs. paracrine signaling, and *N* describes ligand-independent activity (which is the same for every cell, with *N* = 0 in WT and *N* > 0 in the mutant).

The standard parameters of our model, including the parameters that control its WT dynamics, as well as parameters representing several signaling mutants, e.g., the EGF hypomorph *lin-3(e1417)* [24], were previously fit to published data [7], and the same parameter values are used here.

### Quantitative Characterization of Temperature-Dependent Phenotypes for Vulval Patterning

To implement model-guided dynamic perturbations of vulval patterning, we used three temperature-sensitive (ts) signaling mutants; two to manipulate EGF, *lin-15(n765)* and *sos-1(cs41)*, and one to manipulate Notch activity, *lin-12/Notch(n302)* (see Fig. 1F). LIN-15 transcriptionally represses *lin-3*/EGF in the hypodermis, limiting *lin-3* expression to the AC [25] in the WT. In the *lin-15*(*n765)* hypomorphic allele, LIN-3 levels are elevated throughout the hypodermal tissue, leading to ectopic induction of vulval cell fates. At 25°C, all VPCs are induced in this mutant. *sos-1*(*cs41)* is a hypomorphic allele of the Guanine Nucleotide Exchange Factor SOS-1, required for RAS-mediated signaling [26]. The *sos-1*(*cs41)* allele is WT at 15°C and shows strongly reduced induction at higher temperatures (vulvaless, vul). *lin-12*(*n302)* is a weak gain-of-function mutation of the LIN-12/Notch receptor that results in ligand-independent activity [27–29]. *lin-12(n302)* is partially penetrant at both temperatures considered here, 15°C and 25°C.

To quantify the phenotypes of these three mutants and crosses among them, VPC fate patterns at two different temperatures (15°C and 25°C) were scored based on progeny number, cell positions, and morphology of vulval invaginations in the early/mid L4 stage (Fig. 1E). To aid with cell identification and progeny counting, two reporter genes, one labeling the AC and the other labeling VPCs and their progeny, were crossed into the mutant strains (Fig. 1A; we verified that these reporter transgenes do not affect EGF or Notch activity by scoring VPC fate patterns in a sensitized background with reduced EGF, with and without the reporter genes, see Suppl. Fig. 1). Wherever cell fates could not be unambiguously identified, we scored, and the model was fit to, induced (1°/2° fate) vs. non-induced (3° fate) cells.

#### EGF mutants at 15°C

We first considered the *lin-15(n765)* mutant, in which EGF is ectopically produced (Fig. 1F). Contrary to what has been previously reported [21], we found that *lin-15(n765)* is partially penetrant at 15°C (Fig. 2A). Absent any indication to the contrary, we had previously assumed a spatially uniform level of ectopic EGF when modeling that mutant [6,7], and expected comparable induction levels for P3.p, P4.p, and P8.p (Fig. 2B, left-hand panel). However, our observations showed a substantially higher induction of the anterior cells, P3.p (54%) and P4.p (66%), compared to the posterior cell, P8.p (20%). We first considered the possibility that this asymmetry results from a defect in the relative positioning of the VPCs and AC, which has previously been reported to depend on EGF from the AC [30]. Based on the model, a shift in AC position was sufficient to bias P4.p vs. P8.p induction, but P3.p induction remained negligible (Suppl. Fig. 2). Instead, our observations suggested that ectopic EGF levels in *lin-15(n765)* must be non-uniform. Assuming for simplicity a linear gradient of ectopic EGF, parameterized by the offset *δl* and slope *δL* in Eq. (2), we found that this was sufficient to recapitulate the experimentally observed phenotype, with *δL* ≈ 0.18 and *δl* = 0.019 (Fig. 2B-E, Suppl. Movie 3).

**Figure 2:**
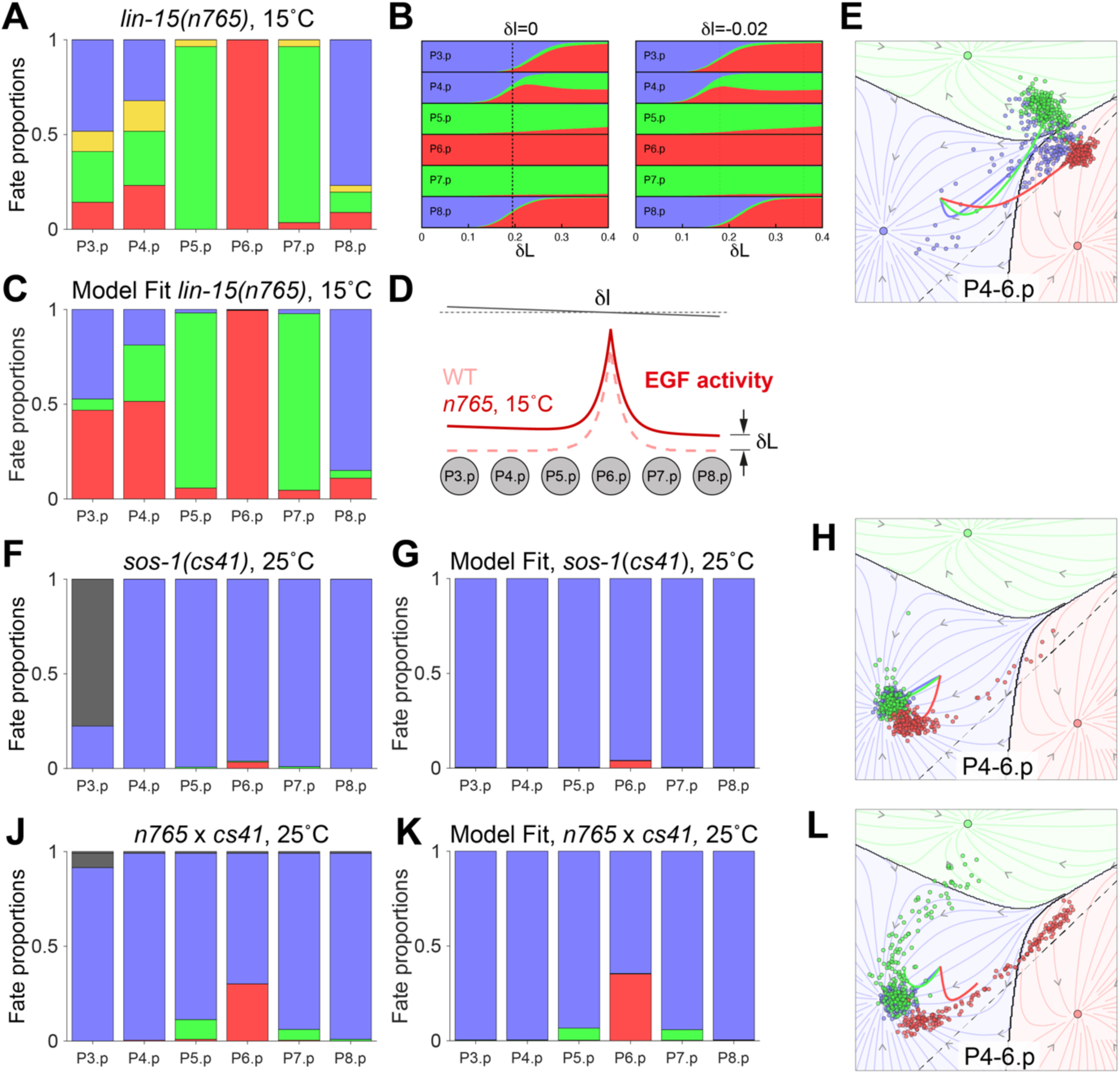
Quantitative analysis and parametrization of temperature-sensitive mutant alleles affecting EGF signaling. (**A**) Fate proportions in the *lin-15*(*n765*) mutant at 15°C (N=27 animals). Yellow denotes induced cells that could not be resolved into 1° vs 2°. (**B**) Fate proportions in the model as a function of ectopic EGF level *δL*, with (right, *δl* = −0.02) or without (left, *δl* = 0) an A-P gradient in ectopic EGF levels, showing the sharpness of the parameter fit and correlations between cells. Dashed lines show model fits to data from **A** (and for the same allele at 25°C in right-hand panel). Parameters are defined in Eq. (1). (**C-E**) Illustration of model fit to the *lin-15*(*n765*) mutant at 15°C, showing the fate proportions (**C**), EGF gradient (solid black line depicts the linear non-uniformity of ectopic EGF) (**D**), and fate trajectories of P4-6.p (**E**; average trajectories and individual outcomes colored according to the WT fate of each cell, cf. Fig. 1E, 200 realizations). (**F**-**H**) Experimental fate proportions (**F**; black filling indicates animals in which P3.p did not become competent), and model fate proportions (**G**) and trajectories (**H**) for the *sos-1*(*cs41*) mutant at 25°C (N=205). (**J**-**L**) Same as (**F**-**H**) for the *lin-15*(*n765*) *x sos-1*(*cs41*) cross at 25°C (N=106). In E,H,L points show cells at the end of the competence period after which they follow the arrows into the valleys for fates 1°-2°-3°,

We then turned to the *sos-1(cs41)* mutant, in which EGF signal transduction is impaired (Fig. 1F). *sos-1(cs41)* on its own is silent at 15°C, and to characterize it we used a cross between *lin-15(n765)* and *sos-1(cs41)*. This showed a slight reduction in induction compared to *lin-15(n765)* alone (Suppl. Fig. 3), from which we infer that the EGF signal transduction is only mildly reduced by the *cs41* mutation at 15°C (fit as *L* ≈ 0.85; this is consistent with the WT phenotype of *sos-1(cs41)* alone, since based on the model, reductions in EGF signaling up to ∼50% are phenotypically silent [6,7], Suppl. Movie 4).

#### EGF mutants at 25°C

At 25°C, induction is almost entirely lost in *sos-1(cs41)* alone, implying a strong reduction of function (fit as a reduction in signaling activity by a factor *L* ≈ 0.18) (Fig. 2F-H, Suppl Movie 5). Complete vulval induction in *lin-15*(*n765*) at 25°C implies a lower bound on EGF levels in terms of the model. To infer a precise model parameter value, we again proceeded indirectly, by analyzing the cross between *lin-15(n765)* and *sos-1(cs41)* (Fig. 2J-L). The cross shows a partial rescue of induction compared to *sos-1*(*cs41*) alone, which we fit to an ectopic EGF level *δL* ≈ 0.35 (Eq. 2) for *lin-15(n765)* corresponding to about twice its value at 15°C; on the other hand, our fits for the slope *δl* at 15°C and 25°C do not significantly differ and we use the same value *δl* = 0.02 in the following. Consistent with the phenotype of *lin-15(n765)* alone at 25°C, the ectopic EGF levels inferred indirectly from the *lin-15(n765) x sos-1(cs41)* phenotype are predicted to be just sufficient for near-complete induction in *lin-15(n765)* alone, as seen in experiments (see the second dashed line in Fig. 2B, right-hand panel, see also Suppl. Movie 6).

#### Notch mutant at 15°C and 25°C

As regards the Notch mutant *lin-12(n302)*, in which the LIN-12 receptor (Fig. 1F) is ectopically activated, animals lack an AC and therefore the source of the LIN-3/EGF inductive signal, due to the role of Notch in the anchor cell vs. vulval uterine (AC/VU) fate decision [13,28,29]. With no AC and constitutive Notch activation, our landscape model predicts roughly equal levels of induction of all VPCs [6]. In contrast, our experiments showed that more posterior VPCs were preferentially induced (Fig. 3A, B). Indeed, the average induction of P5-7.p as a function of temperature was 10% at 15°C and 32% at 25°C, whereas P8.p was only (partially) induced at 25°C, and P3.p and P4.p were never induced at any temperature.

**Figure 3:**
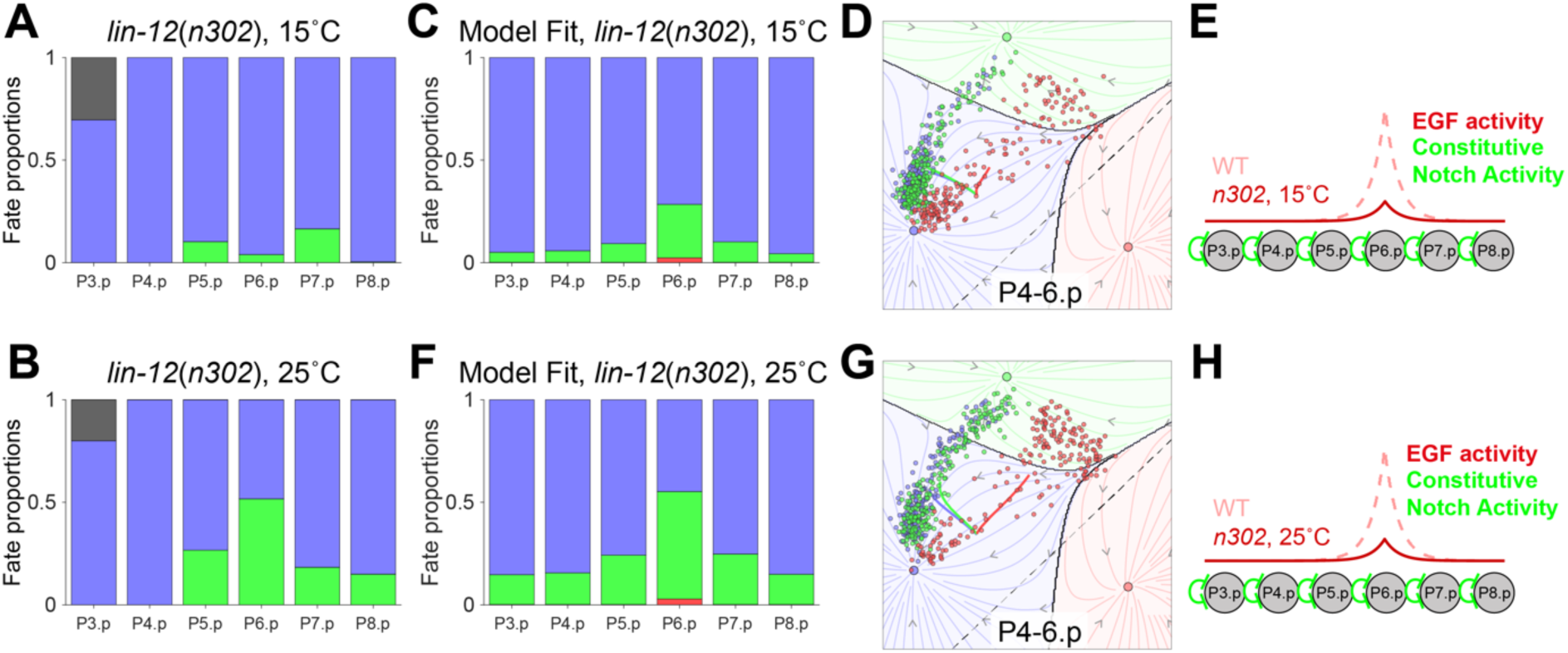
Quantitative analysis and parametrization of a temperature-sensitive Notch mutant allele. (**A**) Fate proportions in the *lin-12*(*n302)* mutant at 15°C (N=112). (**C-E**) Illustration of model fit to the *lin-12*(*n302)* mutant at 15°C, showing the fate proportions (**C**), fate trajectories (**D**), and signaling activity (**E**) including residual EGF and constitutive Notch activity. (**B**,**F**-**H**) Same as (**A,C**-**E**) for the *lin-12*(*n302)* mutant at 25°C (N=30). The fraction of cells of each color that fall into the 3°-fate basins define the partial penetrance.

We reasoned that the higher induction levels of the central VPCs could result from residual *lin-3* expression in the fate-transformed AC, potentially due to incomplete fate transformation. Consistent with this, we occasionally observed expression of GFP from the AC reporter *qyIs362* [lin-29p::GFP] in the somatic gonad of *lin-12(n302)* animals, albeit much weaker than in WT animals (Suppl. Fig. 4). This indicates that the promoter of the AC-specific transcription factor LIN-29, which maintains *lag-2* expression in presumptive ACs, was transiently active for an extended period before cells eventually acquired the VU fate. The anteroposterior gradient in induction levels at 25°C most likely stems from subtle differences in VPC competence present due to low overall EGF activity in this background. VPC competence is established through the combined action of Wnt signaling (via the CWN-1 and EGL-20 ligands) and LIN-3/EGF [31]. EGL-20 forms a long-range anteroposterior gradient across the animal with the source cells in the animal tail [32]. We suspect that, at low EGF levels (due to the absence of an AC), a subtle AP gradient of competence exists that makes posterior cells more sensitive to induction by ligand-independent Notch activity.

To keep model parameters to a minimum, we sought to account for the *lin-12(n302)* phenotype by fitting just two parameters, the prefactor *λ* for the residual EGF gradient from the AC, and the level *N* of ligand-independent Notch activity (in doing so, we ignore the observed A-P asymmetry in experimental outcomes - weak induction of P8.p at 25°C vs. no induction of P3,4.p, as well as the possible time dependence of residual EGF levels from the fate-transformed AC). Based on a fit to our data, both residual EGF and ectopic N activity increase between 15° and 25°C. The changes in the inferred parameter values are relatively small, but being close to the threshold for VPC induction, they are sufficient to result in appreciable changes in induction levels (Fig. 3C-H, Suppl. Movies 7 & 8).

In summary, we have quantitatively characterized the fate patterns of three temperature-sensitive alleles, *lin-15(n765)*, *lin-12(n302)*, and *sos-1(cs41)*, that differentially affect *C. elegans* vulval patterning at various cultivation temperatures. Model fits for permissive temperatures (phenotypically silent or fully penetrant) were achieved through crosses with sensitized backgrounds, employing one of the key strengths of the landscape model, i.e., its ability to predict fate patterns of mutant combinations by simple parameter combination of the individual mutants. The model assumes isotropic time-invariant Gaussian noise in cell-state (see Materials and Methods) and from that noise was able to fit partial penetrance data in all cases by adjusting only deterministic parameters. Thus, the cell-state landscape itself shapes the distribution of outcomes without the need for additional spatiotemporal structure in the cell state noise. This characterization forms the basis for pathway epistasis experiments (crosses between temperature-sensitive alleles and other alleles) as well as time-dependent perturbation experiments, in which we vary the temperature during vulval fate specification to provide time-dependent signaling activities.

### Successful Model Predictions for Epistasis Experiments Provide Insights into Cell Fate Acquisition Dynamics

From the parameter values for three mutant alleles in the EGF and Notch pathways, at 15°C and 25°C, the model now makes direct predictions (free of any additional parameters) for vulval fate patterns for combinations of these alleles with each other as well as other previously characterized mutants in the two pathways [6,7]. As a first test of model predictions, before moving to time-dependent perturbations, we considered different allele combinations under a constant temperature.

#### Epistasis between perturbations within the EGF pathway

Focusing first on mutations in the EGF pathway alone, we considered the effect of ectopic EGF in a background with reduced EGF from the AC, realized experimentally by crossing the *lin-15(n765)* allele into the *lin-3(e1417)* background. *lin-3(e1417)* alone shows about 50% induction of P6.p and very little induction of P5/7.p, a phenotype that was previously fit to a reduction of EGF from the AC by a factor *λ* ≈ 0.28 [7]. To examine the predicted effect of ectopic EGF in this background, we computed the outcome under increasing levels of uniform ectopic EGF (Suppl. Fig. 5A). Strikingly, the model predicted that there is a ‘sweet spot’ just below *δl* = 0.2 where ectopic EGF is sufficient to near fully rescue a WT fate for P5-7.p while causing very little ectopic induction of the more distal cells (P3/4/8.p). This level happens to coincide with our fit for the *lin-15(n765)* allele at 15°C (Fig. 2B), allowing a direct test of the prediction. With allowance for an A-P gradient of ectopic EGF, as inferred in *lin-15(n765)*, the model predicts rescue of P5-7.p and partial ectopic induction of P3/4.p (Fig. 4B,C, Suppl. Movie 9), in remarkable agreement with experimental outcomes (Fig. 4D). Further supporting the model, and the notion that the EGF level at 15°C is marginal for P5-7.p rescue in *lin-15*(*n765*), we construct a triple allele by crossing into *lin-3(e1417)* x *lin-15(n765)* to the *sos-1(cs41)* allele, (previously fit as *L* ≈ 0.85 at 15°C). That results in less complete rescue of P5-7.p, as predicted by the model (Fig. 4E-H, Suppl. Movie 10).

**Figure 4:**
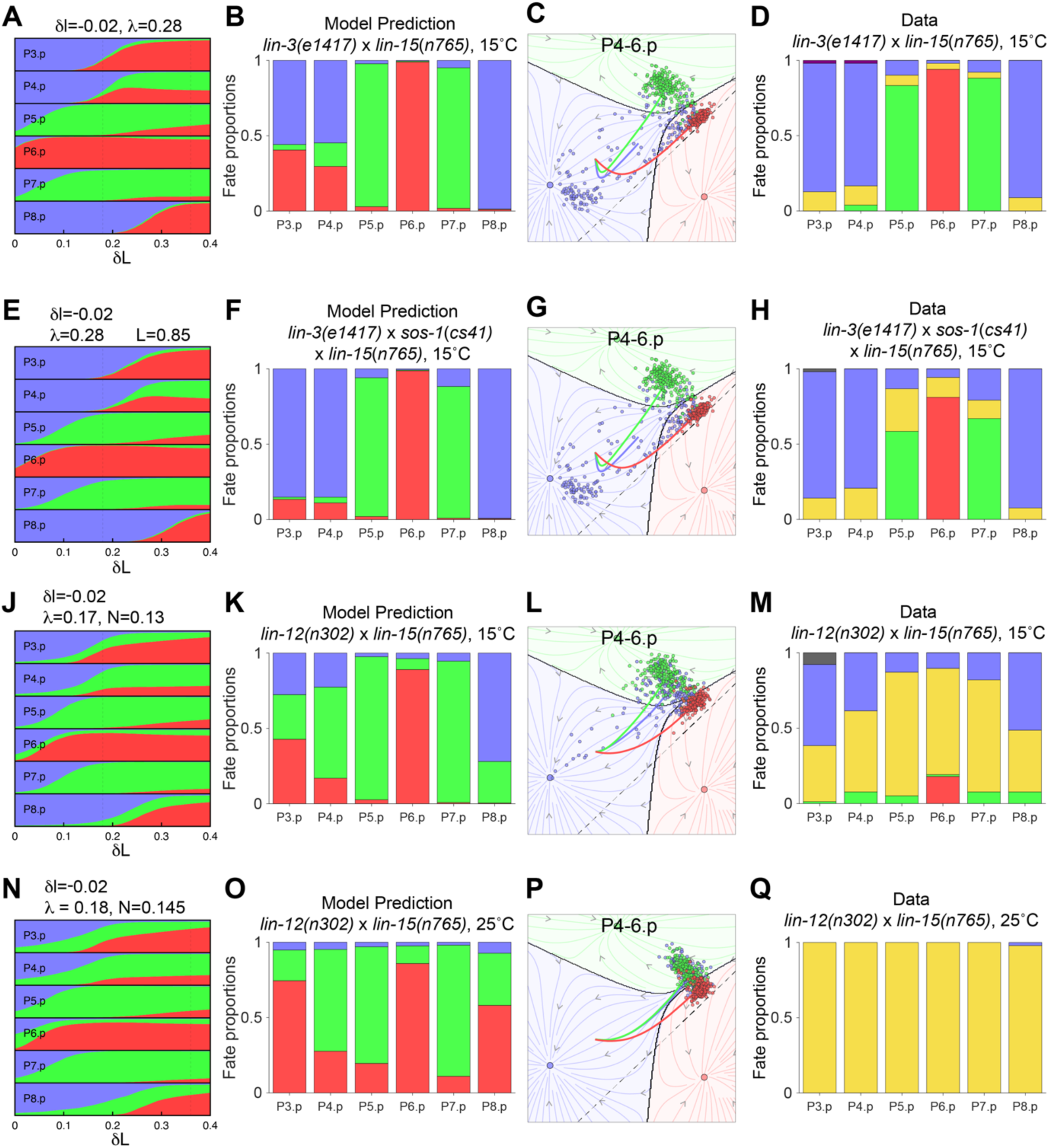
The geometric model quantitatively predicts epistasis of temperature-sensitive alleles. Model predictions are compared with experimental outcomes for crosses between the *lin-15*(*n765*) allele and other mutants within the EGF pathway (**A**-**H**) and in the Notch pathway (**J**-**Q**). In each row, corresponding to a given background and temperature, the first column shows the predicted fate proportions in the model as a function of ectopic EGF level *δL* in that background, with a fixed slope *δl* = −0.02 (**A-D**, *lin-3*(*e1417*); **E**, *sos-1*(*cs41*) *x lin-3*(*e1417*); **J**, **N**, *lin-12*(*n302*)). The following columns show the predicted fate proportions (**B**, **F**, **K**, **O**), predicted fate trajectories (**C**, **G**, **L**, **P**), and experimental fate proportions (**D**, **H**, **M**, **Q**) for the crosses at the indicated temperatures (N=51 for *e1417* x *n765* at 15°C; N=53 for *e1417* x *cs41* x *n765* at 15°C; N=39 for *n302* x *n765* at 15°C; N=24 for *n302* x *n765* at 25°C). The dashed lines in (**A**, **E**, **J**, **N**) show our fits for *lin-15*(*n765*) at the relevant temperature, 15°C or 25°C (*lin-3*(*e1417*) is not temperature-sensitive and the 25°C line is also included in **A** to show the prediction for the cross at that temperature). In experiments, the 1° and 2° fates could not always be distinguished, and some cells were only scored as induced (yellow) or non-induced (blue).

The rescue of P6.p induction to the 1° fate by ectopic EGF is expected, but the rescue of P5/7.p to the 2° fate is less intuitive. Suggesting a possible explanation, it has been found in other contexts that ectopic EGF signaling can synergize with Notch to induce the 2° fate (see [21,6,7]). In our simulations of the *lin-3(e1417)* x *lin-15(n765)* double mutant, however, this effect plays a limited role. Simulations with ectopic EGF restricted to P6.p indicate that the dominant contribution to the induction of P5/7.p is non-cell-autonomous, and results from enhanced Notch signaling from P6.p, which itself is more strongly induced (Suppl. Fig. 6).

#### Epistasis between perturbations in the EGF and Notch pathways

Turning then to conditions in which both EGF and Notch signaling are perturbed, we considered the effect of ectopic EGF in a background with ectopic (ligand-independent and uniform) Notch activity, realized experimentally by a cross between *lin-12(n302)* and *lin-15(n765)*. In *lin-12(n302)* alone, P5-7.p are partially induced to a 2° fate, while the more distal cells are weakly or not induced (Fig. 3; as noted above, we interpret this as reflecting residual EGF signaling in animals lacking an AC). When ectopic EGF is added (here also, we first consider uniform ectopic EGF), low levels of EGF are predicted to result in the rescue of P5-7.p induction and a partial induction of more distal cells to the 2° fate (Suppl. Fig. 5B; by contrast with the previous cross, *lin-3(e1417)* x *lin-15(n765)*, Suppl. Fig. 5A, no level of ectopic EGF is predicted to restore a WT pattern). In terms of the model, the enhanced induction of P5-7.p (which receive low residual EGF in the Notch mutant background) is similar to the rescue of the *lin-3(e1417)* mutant by ectopic EGF, while ectopic 2° fates in more distal VPCs manifest the above-mentioned synergy between low EGF and Notch in the induction of the 2° fate (which in the model arises from the interplay between cell-cell signaling and the nonlinear dynamics of the cells, see [7]). If we allow for a gradient in ectopic EGF as fit for *lin-15(n765)*, at 15°C P3/4.p are predicted to be more strongly induced and yield a mixture of 1° and 2° fates, whereas P8.p is more weakly induced (Fig. 4J-L, Suppl. Movie 11). Although the experimental data do not show as marked an A-P asymmetry, they otherwise agree with model predictions, showing close to full induction of P5-7.p and partial induction in more distal cells (Fig. 4M; the difference between central and distal cells is as expected from the residual EGF in animals without an AC). At 25°C, on the other hand, the ectopic EGF level from *lin-15(n765)* alone is sufficient to induce VPCs, and both model and experiments show near-complete induction in the cross (Fig. 4N-Q, Suppl. Movie 12).

In summary, the two crosses considered here, with a marked difference in the induction of distal cells between 15° and 25°C, thanks to the marked increase in ectopic EGF level (the variation in ectopic Notch activity in *lin-12(n302)* is smaller), lend themselves to dynamic perturbations through temperature shifts, and we can use the model, which correctly captures the outcomes at constant temperatures, to guide the choice of interesting perturbations.

### Guiding cell fates using dynamic perturbations of signaling

So far, we have validated the model’s predictions in terms of static conditions in the genetic backgrounds and end-point phenotypes. As pointed out in [7], time-dependent perturbations of signaling activity during the competence period of precursors provide a much more sensitive test of how well geometric models can capture fate transition dynamics during development. In the model, the time course of signaling can rapidly tilt the epigenetic landscape, thereby determining the cell trajectories. Importantly, signaling dynamics, and thus fate outcomes, can dramatically differ between conditions with identical integrated signaling activities, depending on the timing of this diversion [7]. Viewed from the converse standpoint, geometric modeling can quantitatively predict how specific time courses of signaling may dynamically guide cells towards increased or reduced induction levels or even specific fates.

The temperature-sensitive backgrounds characterized above provide an ideal means to test such dynamical predictions *in vivo*. To do so, our approach was to shift populations of animals with temperature-sensitive mutations at various timings and for various durations during competence from their permissive to their restrictive temperatures (or vice versa). Since for each temperature, we have obtained parameter fits, the model makes quantitative predictions for vulval fate patterns/induction levels as a function of duration and timing of the temperature shifts without additional parameter fitting and making very few assumptions (e.g., immediate parameter change and change of developmental speed upon temperature shift, see Materials & Methods for details).

In the following, we focus on the induction of distal cells (P3/4/8.p), which as noted above (see Fig. 4) show the greatest variation between temperatures: in the two crosses, *lin-12(n302) x lin-15(n765)* and *lin-3(e1417) x lin-15(n765)*, P5-7.p mostly adopt WT fates regardless of temperature, whereas higher ectopic EGF yields higher induction of the more distal cells at 25°C compared to 15°C.

#### EGF pulses in an EGF hypomorph

In a cross with ectopic EGF in a background with reduced EGF from the AC, *lin-3(e1417) x lin-15(n765*), the model predicts that an early pulse of high ectopic EGF (an early 25°C pulse) is much more effective than a late pulse at enhancing the induction of distal cells (Fig. 5A-C and Suppl. Fig. 7A,B, Suppl. Movie 13). Consider first P8.p, that is making a binary decision between the 3°/1° fates. Under low EGF (corresponding to 15°C) the initial state of P8.p generally lies in the basin of attraction of the 3° fate (see first panels in second and third rows in Fig. 5C) so would assume that fate if nothing changes. Under high EGF (corresponding to 25°C), the 3° attractor disappears (undergoing a saddle-node bifurcation with the 3°/1° saddle; see ‘Pulse On’ panels in Fig. 5C), such that all cells would flow into the 1° attractor for constant EGF. Thus, the duration of an early EGF pulse determines the fraction of cells that traverse the 3°/1° saddle and go to 1° while with a late pulse, cells sink further into the 3° basin and cannot be kicked out by EGF. As depicted in Fig. 7, this essentially cell-autonomous effect is recovered in a simpler, 1D model for the decision of an isolated cell between two fates. As a slight departure from this simple picture, we note that P8.p occasionally adopts the 2° fate following an early pulse (in both model and experiments; Fig. 5B-D), which can be attributed to the higher induction of P7.p to the 1° fate and resulting Notch signaling to P8.p.

**Figure 5:**
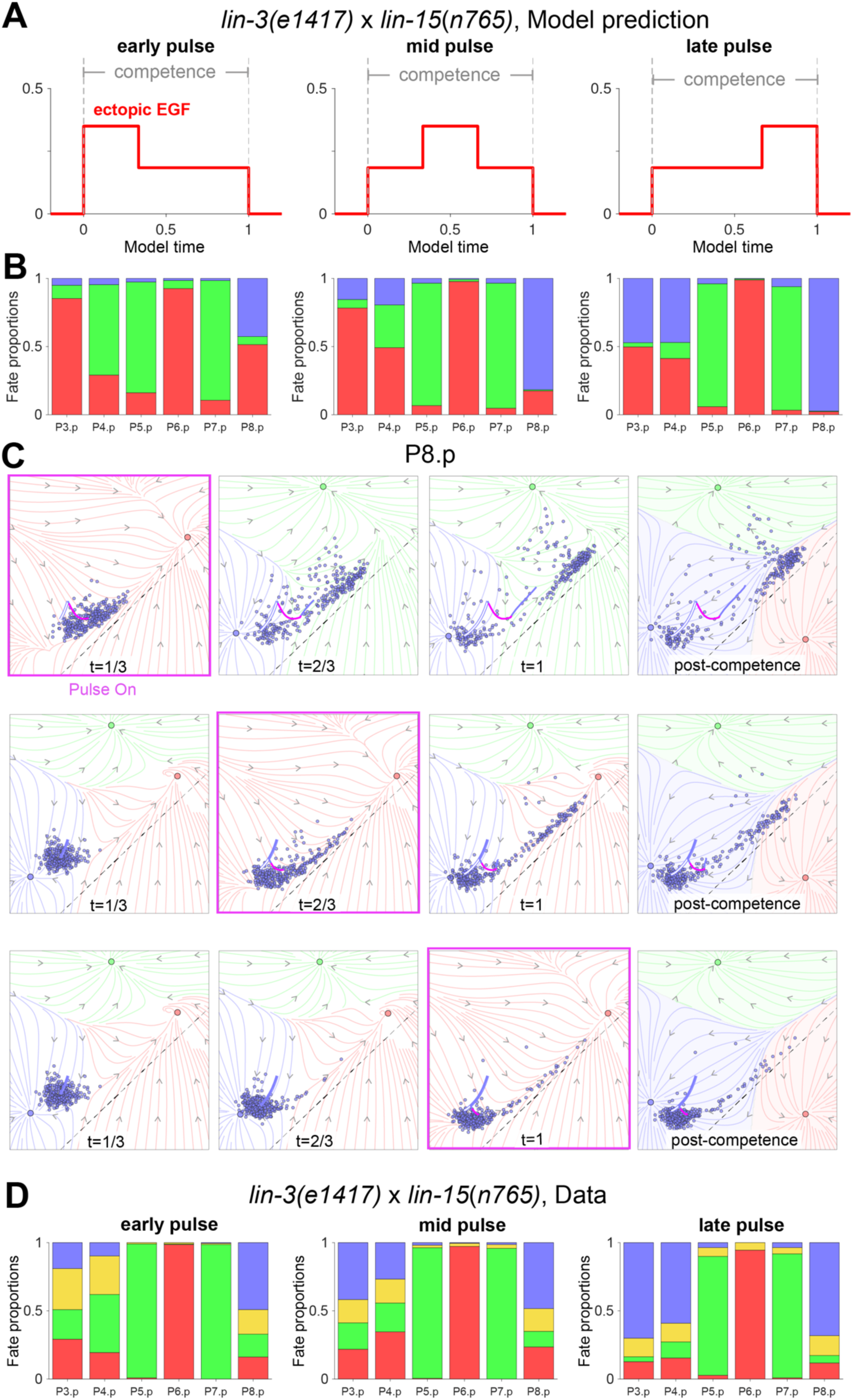
Quantitative guiding of VPC fates with timed pulses of signaling activity in a background with reduced EGF from the AC. (**A**) Temperature shifts at different times during competence in the *lin-3(e1417)* x *lin-15(n765)* double mutant are modeled as step-like increases and decreases in ectopic EGF, with levels based on our fits for *lin-15(n765)* at 15°C and 25°C. (**B**) Predicted fate proportions of P8.p for early (left), mid (middle) and late (right) temperature shifts. (**C**) Predicted fate trajectories for P8.p for early (top row), mid (middle row) and late (bottom row) temperature shift, based on our fits reduced EGF from the AC in *lin-3(e1417*) and graded ectopic EGF in *lin-15*(*n765*). To show how the time-dependent dynamics shape the trajectories, we depart from previous figures by overlaying for successive time intervals of 1/3 of the competence period (ending at the indicated time points) the fate trajectories on flow fields that integrate the EGF and Notch signals received by P8.p (paracrine Notch signaling is averaged over each time interval and all realizations [N=300]; see Methods). In each panel, the portion of the average trajectory corresponding to the pulse is highlighted in magenta, and the average trajectory in the absence of a pulse is shown as a white curve with a blue outline. The last column shows the flow field without signals, with determines cell fates post competence. This representation illustrates our interpretation of the outcomes based on the position of the cells at the end of the pulse and dynamics after the downshift, e.g., an early pulse pushes (on average) P8.p to the separatrix (interface between blue trajectories ending in 3° and red trajectories ending in 1°) that exists at 15°C, such that cells subsequently go to a mixture of 3° and 1° fates. (**D**) Experimental fate proportions for early (left, N=108), mid (middle, N=112) and late (right, N=56).

The situation is similar for P3,4.p, except that they receive a higher EGF level (and therefore show stronger induction even at 15°C), and Notch signaling between P3,4.p results in induction of 2° as well as 1° fates (this is especially the case for P4.p, see Suppl. Fig. 7). Here also, the same integrated dose of signal is predicted to displace fate outcomes much more effectively when delivered early vs. late. It is noteworthy that the flow fields for P4.p (Suppl. Fig. 7) show a ‘decision structure’ that is largely similar to P8.p (Fig. 5C), with trajectories spreading along the direction between the 3° and 1° fates (see the low-EGF flow field in the first panels of the second and third row in Suppl. Fig. 7, and notice the cells near the saddle between 3° and 1°), with the difference that the trajectories of induced cells eventually curve towards the 2° fate under the influence of Notch signaling. In other words, this timed perturbation transforms the normal three-way cell fate decision structure (in which the trajectories of presumptive 2° cells cross a saddle between the 3° and 2° fates, Fig. 1E) into one that is comprised of two sequential binary fate decisions: cells first ‘decide’ between non-vulval (3°) and vulval (1°/2°), then following the high EGF pulse, among the induced cells, the balance between the EGF and Notch (which depends on cell position) governs which go to a 1° or 2° fate.

The predictions of the model are fully borne by experiments in which animals were shifted from 15°C to 25°C and back at different times during competence (Fig. 5D and Suppl. Fig. 8A). Importantly, to rule out the possibility that later pulses were less effective because they extended beyond the period when VPCs are responsive, we ensured that the pulses lay within the competence period, based on the timing of the first VPC divisions (see Materials & Methods). Beyond the higher induction levels for early vs. late pulses, the model also predicts a higher fraction of 2° cells among the induced cells for P8.p following an early pulse (Fig. 5B), which can be attributed to the longer time during which Notch signal can act on induced cells after the pulse (Fig. 5C). In experiments, a 1° vs. 2° fate could not be assigned to all induced cells, but the data for P8.p when a fate was assigned indeed show a higher 2° fraction following early pulses (early [N=110 animals] vs. mid [N=108]: p < 0.003; early vs Late [N=55]: p < 0.04; mid vs. late: p < 0.3715; early vs. mid and late [pooled]: p < 0.0003), providing further support to our interpretation of the response to dynamic perturbations.

#### EGF pulses in a Notch mutant

In the case of ectopic EGF in a background with low ectopic Notch, *lin-12(n302) x lin-15(n765*), the model similarly predicts that an early pulse is much more effective than a late pulse at displacing the outcomes [7]. Specifically, an early pulse of high EGF can result in near-complete induction of P3,4.p and P8.p, and the increase of induction compared to constant low EGF is mostly in the form of 2° fates (compare Figs. 4K and 6A,B), at odds with the suggestion that EGF will promote the 2° fate specifically at low levels [21] (in which case a pulse of high EGF would be expected to induce 1° fates). Here also, as in the above case of EGF pulses in an EGF hypomorph, the decision structure is one in which the cells first decide between non-vulval and vulval, before Notch (which here dominates over EGF) diverts the induced cells to the 2° fate. As a noteworthy consequence of this, the predicted outcome of an early pulse in this case markedly differs from that of a constant high EGF level, in which ectopic 1° fates predominate (compare Figs. 4O and 6B).

**Figure 6:**
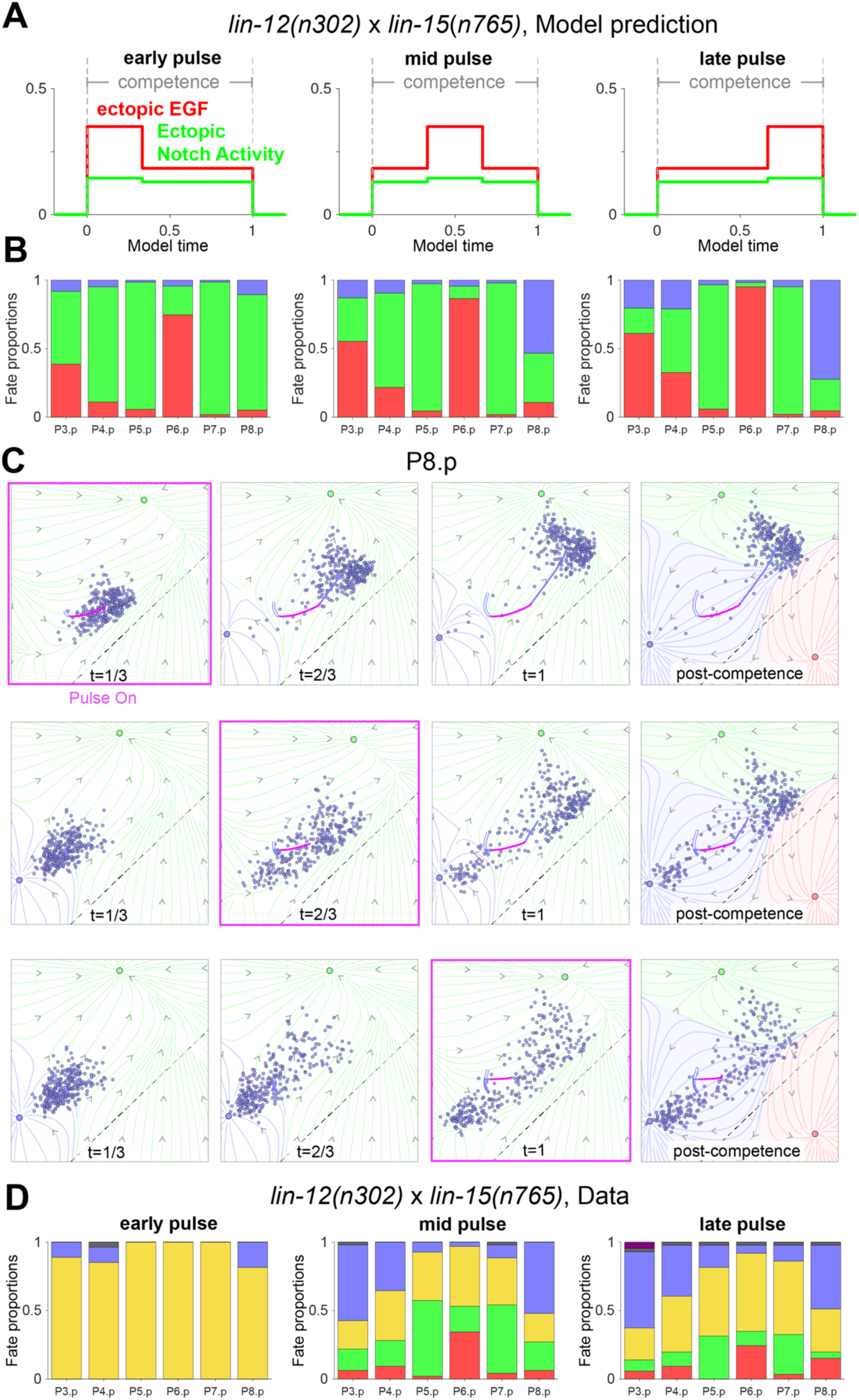
Quantitative guiding of VPC fates with timed pulses of signaling activity in a background with reduced EGF from the AC. (**A**) Temperature shifts at different times during competence in the *lin-12(n302)* x *lin-15(n765)* double mutant are modeled as step-like increases and decreases in signaling activities, based on our fits for the two alleles at 15°C and 25°C. (**B**) Predicted fate proportions of P8.p for early (left), mid (middle) and late (right) temperature shifts. (**C**) Predicted fate trajectories for P8.p for early (top row), mid (middle row) and late (bottom row) temperature shifts. Flow fields determined as described in Fig. 5C, but based on fits for residual EGF from the AC and constitutive Notch activity in *lin-12(n302)*, and ectopic EGF in *lin-15(n765)*. The portion of the average trajectory corresponding to the pulse is highlighted in magenta, and the average trajectory in the absence of a pulse is shown as a white curve with a blue outline. (**D**) Experimental fate proportions for early (left, N=30), mid (middle, N=42) and late (right, N=38).

**Figure 7:**
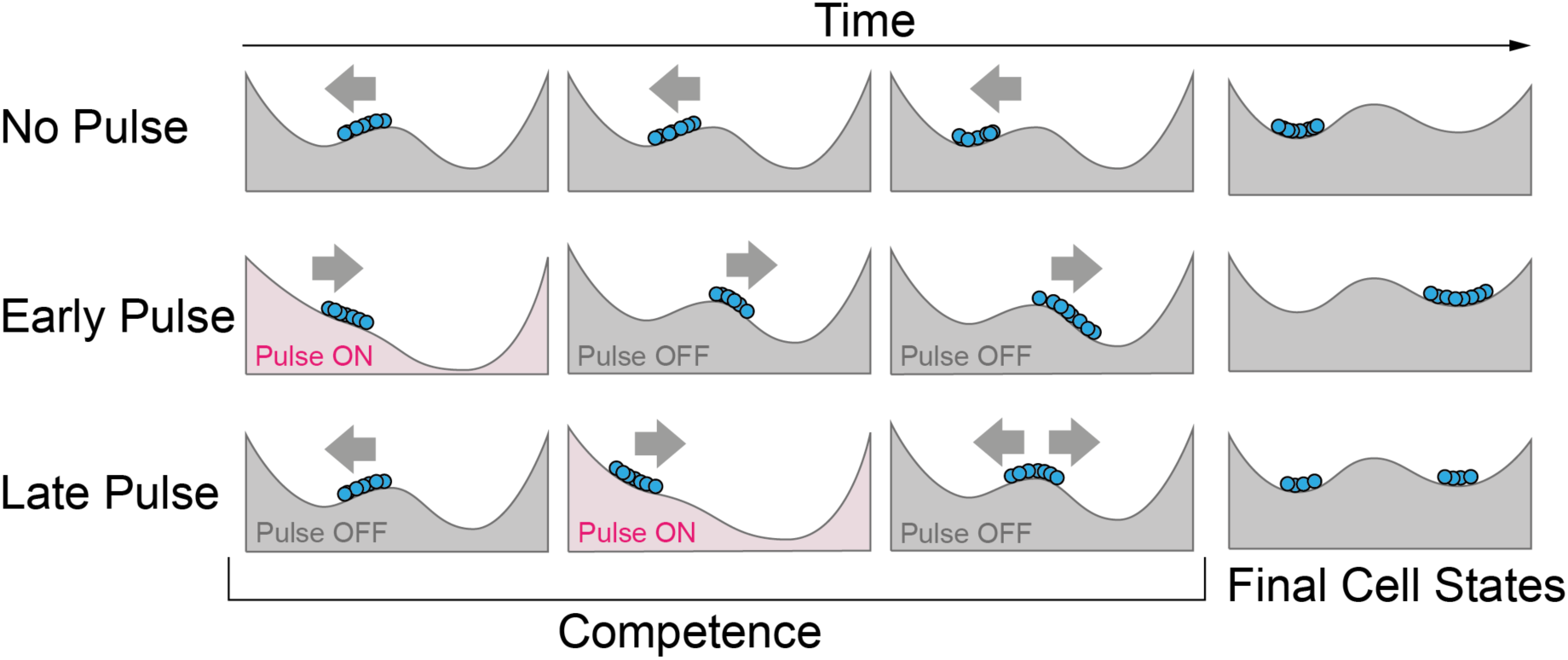
A mechanism for dynamical guidance of cell fates in a Waddington Landscape through pulses of signaling. Without a pulse, cells move to the left valley. Early pulsing pushes cells far enough to the right to convert the fate of all cells. A pulse later in the competence period does not convert fates as efficiently.

This further argues against the notion of attaching a unique fate to EGF signaling. Consistent with our predictions, experiments in which animals were shifted from 15°C to 25°C and back at different times during competence here also showed a much greater induction by early vs. late pulses (Fig. 6D, Suppl. Fig. 8B, Suppl. Fig. 9, and Suppl. Movie 14).

Together, our temperature shift/signaling pulse experiments provide a quantitative *in vivo* test of a landscape model with dynamic perturbations to cell signaling, both within a single signaling pathway and involving pathway epistasis. They also show how this modeling approach can provide intuitive explanations for transition dynamics of cell fate patterns that inform strategies for how these transitions can be optimized by controlling signal timing.

## Discussion

Recent years have seen a surge of interest in giving the Waddington landscape a mathematical foundation to test its predictive value with quantitative experimental data. By now, several studies have shown that geometric or landscape models, which abstract away much of the often system-specific molecular details, provide a general framework to systematically classify possible scenarios for cellular decisions [6,7,9,33], and are remarkably adept at capturing the essential features of cellular decision making in concrete systems [8,12]. They are rich enough in parameters to allow specific attribution of the alleles we studied to parameters. Apart from *in vitro* experiments in ‘decoupled’ mESCs (with no intercellular signaling) [12], experimental and theoretical studies have so far focused on endpoint descriptions and static perturbations (mutant cell lines/strains). Here, using classic temperature-sensitive alleles of the signaling pathways involved in *C. elegans* vulval patterning, we developed an approach to directly demonstrate that geometric models can capture dynamic cell-cell interactions, which are very relevant in the context of spatial patterning *in vivo*, but also generically occur through ligand production and intrinsic signaling between initially equivalent stem cells *in vitro*. We have shown that a previously developed landscape model can capture vulval fate patterns for several temperature-sensitive alleles of the EGF/Ras/MAPK and Notch pathway when raised at either their permissive or restrictive temperatures. Model fits for phenotypically silent conditions could be obtained by fitting crosses with sensitized genetic backgrounds, highlighting one of the key predictive strengths of the model. When we then performed dynamic perturbation experiments, by shifting populations of animals from permissive to restrictive and back to permissive temperatures during precursor competence, the model correctly anticipated the observed dependencies of fate patterns on the timing and duration of the temperature shift (i.e., of the signaling ‘pulse’). Our results demonstrate a general approach that leverages interpretable models to predictively guide individual cell fates and collective fate patterns, with obvious implications for the rational engineering of tissues and organs, e.g., for regenerative medicine.

Beyond challenging model predictions against the eventual fate patterns, the predicted dynamics could in many contexts be directly compared with data from reporters. In the context of vulval patterning, several reporters have been used previously to identify VPC fates, but they do not systematically correlate with fate in partially penetrant backgrounds [34], making it unclear that they provide a faithful readout of developmental trajectories. On the other hand, live sensors for the ERK and N pathways have recently been developed, that could enable more dynamic experiments, while keeping in mind the distinction between the pathway *activity* that we model from the nuclear translocation of pathway effectors that is probed with these reporters. For the EGF pathway, the activity of ERK, the most downstream protein kinase of the canonical Ras-Raf-MEK-ERK cascade, has been monitored via a translocation reporter [35]. As in other model systems [36–38], ERK activity was found to be pulsatile, with pulse frequency increasing with increasing ligand levels. How this relates to upstream pathway *activity*, which our model purports to describe, and is interpreted in the expression of the receiving genes, remains to be worked out. For Notch activity, a biosensor called SALSA based on an endogenous tag of LIN-12 with a TEV protease and co-expression of GFP-(TEV-cut-site)-RFP has recently been published [39]. One key observation from this biosensor was the consistently higher sensor signal in P5/7.p vs. P6.p, when our model predicts the highest Notch activity for P6.p. throughout competence (due to the high ratio of autocrine to paracrine signaling, as required to account for the adoption of the 2° fate by isolated cells upon ablation of all but one VPC [40–42]). Besides the ambiguities in terms of the exact measure of pathway activity readout by the SALSA Notch sensor, we speculate that elevated Notch activities in P5/7.p compared to P6.p may not be required for fate patterning but rather features of fate execution. Indeed, a variation of our model that explicitly incorporated a downregulation of Notch signaling in presumptive 1° cells allowed an eventual shutoff of autocrine signaling, and a closer approach to the 1° fate attractor by those cells, but otherwise left the dynamics of patterning and its response to perturbations largely unchanged [7].

Genetic screens over the past 40 years have likely uncovered a near-complete molecular parts list underlying the specification and execution of VPC fates. With few exceptions, these discoveries have been made by scoring endpoint fate phenotypes and clever genetic reasoning. Live imaging and perturbations have seldom been used in this system, due to the technical challenges of observing feeding, growing, and moving animals for extended periods at high resolution. Recent advances in microfluidics for developing *C. elegans* larvae [43,44] should render long-term observations and live manipulations during imaging feasible and promise to open new avenues for quantitative studies of cell fate acquisition dynamics. We anticipate that, through combinations of these imaging techniques with tools for real time manipulations of signaling, e.g., via optogenetic control of protein localization [45–47], *C. elegans* vulval development will continue to yield increasingly quantitative insights into *in vivo* cellular differentiation.

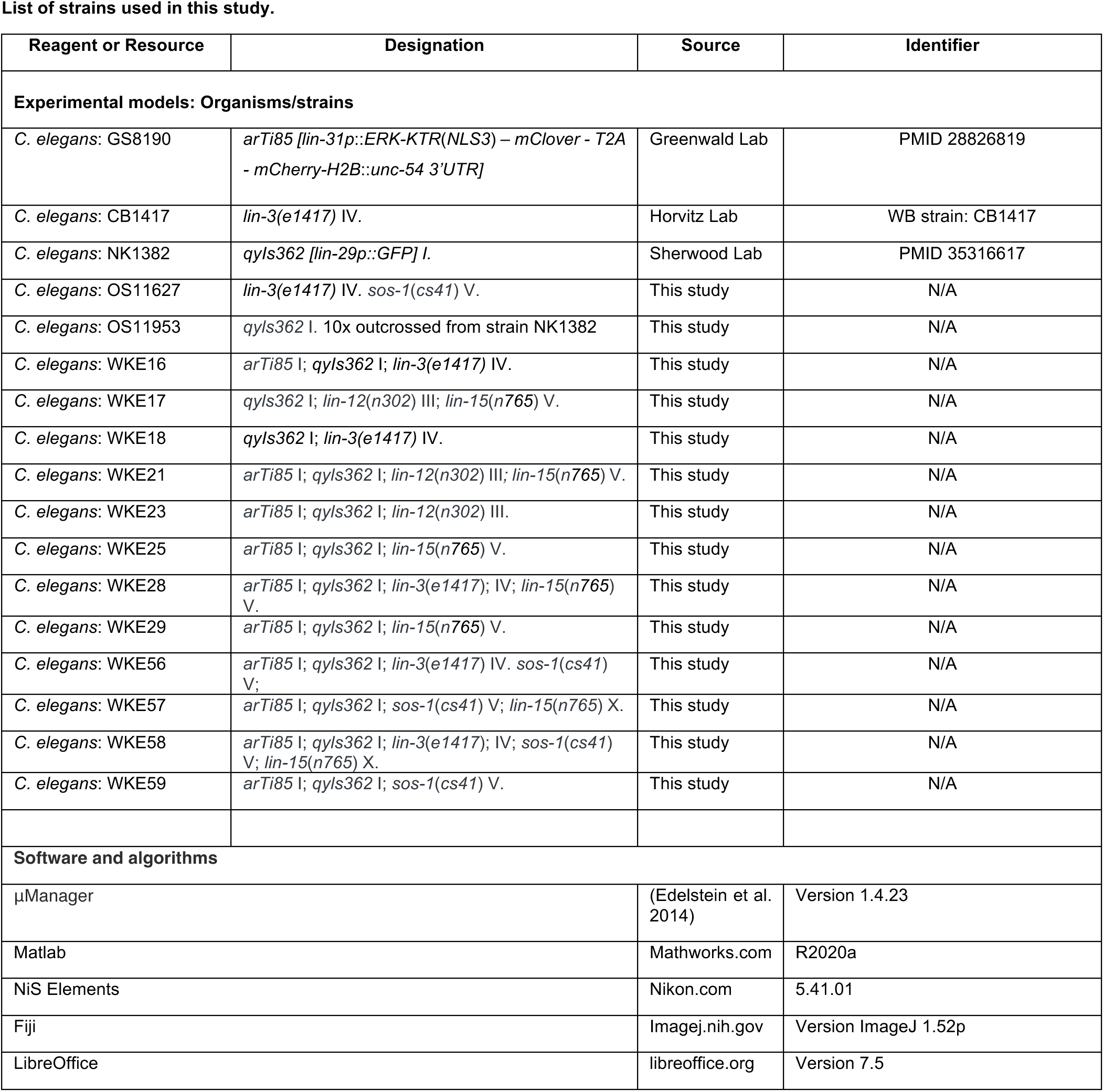

## Materials and Methods

### C. elegans strains and general culture conditions

A complete list of strains used in this study is given in Table 1. *C. elegans* stocks were maintained on 2.5% agar NGM (Nematode Growth Medium) plates (55mm diameter) seeded with *E. coli* strain OP50 at 20°C unless noted otherwise [48]. Mixed-age hermaphrodite stock cultures were hypochlorite-treated to obtain age-synchronized, arrested L1 larval populations [48]. All strains were raised at 15°C and shifted to the indicated cultivation temperature upon feeding after L1 starvation arrest. To avoid the strong larval lethality of *sos-1(cs41)* at elevated temperatures, *sos-1(cs41)* animals were maintained at 15°C for 30h after L1 starvation arrest (mid L2 stage) before being shifted to 25°C.

**Table 1.** Model Fits Summary of fate induction patterns and model parameter fits for EGF/Ras/MAPK and Notch alleles characterized in this study. All alleles, except *cs41* at 15°C, have *arTi85* I; *qyls362* I in the background (see Materials & Methods for details). Fates are temperature-independent in *lin-3(e1417).* In *sos-1(cs41)* at 15°C, the fate pattern is 100% WT and in *lin-15*(*n765)* is fully induced (see Materials & Methods for details).

| Allele | T(°C) | VPC fates<br>(% 1°, 2°, 3°) or (% induced, non-induced) |  |  |  |  |  | Fitted Model<br>Parameters | Reference For Model<br>Fit |
| --- | --- | --- | --- | --- | --- | --- | --- | --- | --- |
|  |  | P3.p | P4.p | P5.p | P6.p | P7.p | P8.p |  |  |
| *lin-15(n765)* | 15°C<br>(N=27) | 54,46 | 67,33 | 100,0 | 100,0 | 100,0 | 20,80 | $\delta L = 0.18$<br>$\delta I = -0.02$ | This study |
| | 25°C | 100,0 | 100,0 | 100,0 | 100,0 | 100,0 | 100,0 | $\delta L = 0.35$<br>$\delta I = -0.02$ | This study |
| *sos-1(cs41)* | 15°C | 0,0,100 | 0,0,100 | 0,100,0 | 100,0,0 | 0,100,0 | 0,0,100 | L = 0.85 | This study |
|  | 25°C<br>(N=205) | 0,0,100 | 0,0,100 | 0,0,100 | 4,0,96 | 0,0,100 | 0,0,100 | L = 0.18 | This study |
| *lin-3(e1417)* | (N=88) | 0,0,100 | 0,0,100 | 0,10,90 | 54,0,46 | 0,10,90 | 0,0,100 | $\lambda = 0.28$ | (Corson & Siggia,<br>2017) |
| *lin-12(n302)* | 15°C<br>(N=112) | 0,0,100 | 0,0,100 | 0,10,90 | 0,4,96 | 0,17,83 | 0,1,99 | N = 0.13<br>$\lambda = 0.17$ | This study |
| | 25°C<br>(N=30) | 0,0,100 | 0,0,100 | 0,27,83 | 0,52,48 | 0,18,82 | 0,15,85 | N = 0.145<br>$\lambda = 0.19$ | This study |

### VPC fate scoring and determination of developmental age

Following convention, we scored animals for VPC fates via Nomarski microscopy after cessation of vulval cell divisions (early/mid forth larval stage (L4). Previous studies used the number of progenies, axis of cell divisions, adherence to the ventral cuticle, overall morphology of the vulval tissue gathered from Nomarski microscopy as hallmarks of fate choice [17,18]. We found cell-fate assignment in *muv* strains based on those criteria highly ambiguous. Therefore, we crossed all characterized alleles into transgenes *arTi85* [lin-31p::ERK::KTR::mClover::T2A::mCherry::his-11::unc-54 3’UTR + rps-27p::NeoR::unc-54 3’UTR] and *qyIs362*[lin-29p::GFP]. *arTi85* includes a histone H2B mCherry tag, expressed in VPCs and its descendants (Fig. 1A). *qyIs362* is an AC-specific cytoplasmic GFP reporter, allowing unambiguous identification of the AC in confocal stacks (Fig. 1A). The strain NK1382, harboring *qyIs362* and *qyIs17 [zmp-1p::mCherry]* II [49], was outcrossed, to obtain our *qyIs362* background WT strain OS11953 (see Table 1). The H2B mCherry tag is also expressed in P1-2.p and P9-11.p nuclei at the L4 stage. P1-2.p, are the first two cells and P9-11.p the three last cells labeled in the ventral mid-line from the pharynx going to the posterior and centered laterally. These cells do not divide and are, thus, bigger than terminally differentiated VPCs, serving as a landmark for identifying P3.p, P4.p, P5.p, P6.p, P7.p, P8.p or their progeny. VPC fates were assigned as follows:

- 1° lineage undergoes two rounds of division to produce four cells that divide transversely.
- 2° lineage undergoes two rounds of division to produce four cells the two external cells divide laterally and adhere to the ventral cuticle; the two inner cells detach from the ventral cuticle the most external one divide transversely and the most inner one does not divide.
- 3° lineage undergoes one round of division to produce two daughters that cease to divide and fuse with the main body epidermis.

In most of the mutants deviating from the WT pattern, VPCs can adopt half fates, i.e., the two daughter cells follow different lineages from 1°, 2° or 3°. Half fates involving 3° are easily identifiable as the 3° daughter cell is bigger than the 1° or 2° progeny. Disambiguation between 2° and 1° was done, when possible, based on the criteria in [41] (adhesion to cuticle, axis of division and symmetry and morphology of the invagination). When no clear distinction was possible, fate was noted Vulval or V. For model fits, half-fates were replaced by two individuals in which the cell fully adopted one of the two fates.

Developmental age of each animal was based on the same images used for scoring vulval fates. The specific developmental substage (L4.0 to L4.9 [50]) was determined based on up to four anatomical criteria: (1) morphology of the vulval invagination [50], (2) molting status (presence of a mouth cap, detached cuticle, used for L3 vs. L4.0 vs. L4.1), (3) Anchor cell morphology (used for L4.1 – L4.4), and (4) extend of migration of the distal tip cell [51] and gonadal elongation. The anchor cell is also used as another landmark, as it is normally positioned directly dorsal to the P6.p progeny.

### Model formulation and implementation

The geometric model [6,7], describes the state of each VPC by a vector 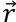 in two-dimensional space (Fig. 1C) and its dynamics by the stochastic differential equation

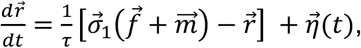

where 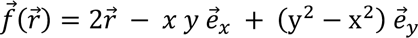 is a polynomial vector field with threefold symmetry and the nonlinear function

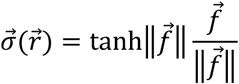

ensures that the dynamics is bounded. The term 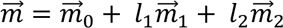 integrates a bias towards the default 3° fate, 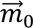, and the effect of EGF and Notch signaling, parameterized by two vectors, 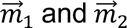, which are combined linearly according to the levels of EGF and Notch ligands, *l*_1_ and *l*_2_ on the cell (see main text). Variability in the dynamics is described by the stochastic term 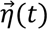, parameterized by a coefficient of diffusion *D* in phase space,

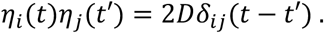

A fixed exponential gradient of EGF is assumed

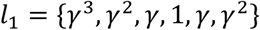

while the level of Notch ligands a cell exposes to its neighbors is a function of its current state

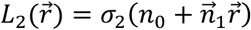

and varies continuously from 0 to 1 across a line parameterized by *n*_0_ and *n* (Fig. 1C, bottom right). Stochastic simulations of the model were performed as described in [6].

### Parameter fitting

All model parameter values were taken from [7], except noted otherwise. Ligand levels were fit to the proportions of the different fates, 1°−3°, adopted by each cell, P3-8.p, in different conditions (correlations between cells are thus ignored; as in [6], the fit minimizes the sum of squares of the difference between fate proportions in model and experiment). For conditions in which we were unable to distinguish between 1° and 2° fates (e.g., *lin-15(n765)*), we fit the fraction that each cell adopted either 1° or 2° fate in the model. As an exception to this general procedure, the model was fit to the *lin-12(n302)* allele at 15°C based on the average fate proportions among two groups of cells, P4,8.p and P5-7.p (fit in this way, the model captures the higher induction of central VPCs but exhibits highest induction in P6.p, at odds with the experimental data, see Fig. 3A,C; when it was fit to the full fate pattern, it failed to capture any difference within the VPCs, yielding uniformly low induction).

#### Time-dependent flow fields integrating EGF and Notch signaling

In Figs. 5 and 6 and Suppl. Figs. 7 and 9, cell trajectories during successive time intervals are overlaid over flow fields integrating the EGF and Notch signals received by the cell. These are defined by the vector field 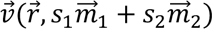 as in Eq. 1, where the EGF signaling activity *s*_1_ of the cell (which is constant within each of the considered time intervals) is taken from Eq. 2, and Eq. 3 is adapted as follows to treat separately the autocrine and paracrine contributions to the Notch signaling activity *s*_2_ of the cell (e.g. P6.p):

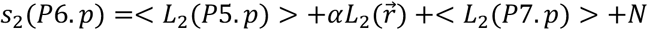

In this equation, *N* denotes as usual ligand-independent activity (which is constant within each time interval). The brackets indicate that the paracrine contributions from P5.p and P7.p, which vary in time and between realizations, are averaged over each time interval and all realizations to define a single flow field that best represents the dynamics of the cell during that interval. By contrast, the time-dependent autocrine contribution *αL*_2_(*P*6. *p*), which depends on the current state of the cell, can be built into the ‘landscape’ in the form of the term 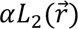, to adequately represent how the trajectories of cells in different states (e.g. for different realizations) are deflected by autocrine Notch signaling.

#### Inferring timing and duration of signaling perturbations via temperature shifts

Inferring the timing and duration of signaling perturbations via temperature shifts requires an estimation for how much “Developmental time” has passed in each absolute time and, ultimately, what fraction of competence was spent at which temperature. In the following, upper case times indicate “Developmental time” in **model time units**, lower case times indicate **absolute** times in hours. We define:

- *v*_0_ … developmental speed at 15°C, measured in “(developmental time)/h”.
- *α* … dimensionless scaling factor for developmental pace at 25°C vs. 15°C, so that *v*_25°*C*_ = *αv*_0_. For WT *C. elegans* animals, *α* can be inferred comes from the Arrhenius law for developmental rate of *C. elegans* (37). For other genotypes, e.g., *lin-3(e1417)* x *lin-15(n765)*, we measured developmental progression at 25°C vs. 15°C through staging age-synchronized populations at various absolute times of development in the L3 and L4 stages. *α* was found to be ∼1.4 for all strains.
- *a* … acceleration in developmental speed after temperature switch, measured in “(developmental speed)/h”. We assume that acceleration is linear and *α* = *γv*_0_ with *γ* in units of 1/h. This formulation assumes that all worms take the same **absolute** time to increase their own developmental speed by a certain factor, i.e., faster developing worms accelerate at a faster absolute rate.

We define the absolute times *t*_1_, *t*_2_, *t*_3_, *t*_4_ as the time spent at 15°C, time spent accelerating to 25°C developmental rate, time spent at 25°C, and time at 15°C developmental rate (assuming instantaneous deceleration), the total developmental time passed when the animal is scored is given by

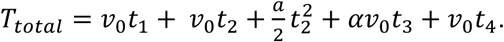

From the experiment, known are the absolute times *t*_1_, *t*_*pulse*_ = *t*_2_ + *t*_3_, and *t*_4_. With the assumptions above, *t*_2_ = *α*/*γ*. Solving the equation for *T*_*total*_ for *v*_0_gives the developmental time (model time) passed per hour at 15°C. From this, one can calculate how much developmental time has passed in all temperature phases of the experiment.

In line with prior work, we assumed VPC competence to last from the mid L2 stage until shortlt after the first VPC division, which we determined to be at 1/3 in the L3 stage in WT, *sos-1(cs41)* and *lin-3(e1417)* (data not shown). This developmental time span corresponds to the model time interval [0,1]. In line with recent studies on temporal scaling in *C. elegans* post-embryonic development [52], we assumed that all animals have the same relative duration of each larval stage, independent of their individual developmental pace. Furthermore, we assumed that animals immediately shift developmental pace upon temperature shifts (i.e., *t*_2_ = 0). To ensure that the late temperature pulses were delivered within the VPC competence window, we verified in a subset of the populations that, at the time points of the temperature down shifts, VPCs had not undergone their first round of division in the vast majority of animals.

## Data availability

All strains used in this study (Table 1) are available upon request.

## Acknowledgments

We thank Eric D. Siggia for guidance and advice throughout all project stages. We thank David Sherwood for sharing the NK1382 strain ahead of publication. We thank Rémy Fert and Giacomo Gropplero (Institut Curie) for help in building a custom temperature-control system on our microscope. We thank Shai Shaham, members of Shaham lab (The Rockefeller University) for feedback during the early stages of this project. We thank Marie-Anne Félix (ENS, Paris) for advice on VPC fate scoring. We are grateful to all members of the Keil lab for fruitful discussions and feedback throughout this project. Some strains were provided by the *Caenorhabditis* Genetics Center (CGC), which is funded by the National Institute of Health Office of Research Infrastructure Programs (P40 OD010440).

## Author contributions

FC and WK conceived and designed the project. IH and WK performed all experiments, supervised by WK. IH, FC, and WK analyzed the data. All authors wrote and edited the manuscript.

## Supplementary Figures and Figure Legends

**Supplementary Figure 1:**
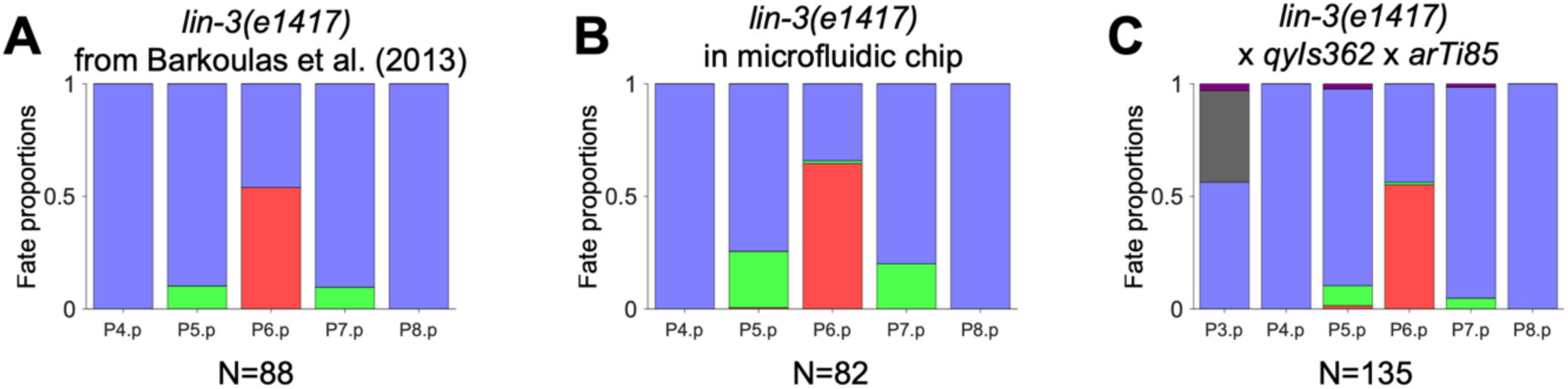
The AC marker *qyls362* and the ERK biosensor *arTi85* do not influence VPCs fate decisions. To assert whether AC marker *qyls362* and ERK biosensor *arTi85* affect either EGF or Notch pathway activity in VPCs, we crossed these transgenes into the *lin-3*(*e1417*) sensitized background. The allele was previously fitted as *λ* = 0.28 (see Eq. (2)) based on data from [53] (**A**). We observed almost identical fate proportions for *e1417* when scored from animals developing in a previously published microfluidic device [43], showing that *e1417* is robust with respect to environmental conditions (see also [53,54]) (**B**). Any perturbation in the EGF or Notch signaling pathways are predicted to dramatically change the fate proportions as changes in *l*_1_ or *l*_2_ (representing EGF and Notch ligand levels) by only 0.05, for example, would lead to either 0% P6.p induction or double the induction levels of P5/7.p. However, when crossed into the two fluorescent reporters (**C**), parameter fits placed this strain within the confidence intervals of *e1417*. This is consistent with the hypothesis that EGF emission from the AC and MAPK & Notch pathway activities in VPCs are not affected by the *arTi85* and *qyls362* transgenes.

**Supplementary Figure 2:**
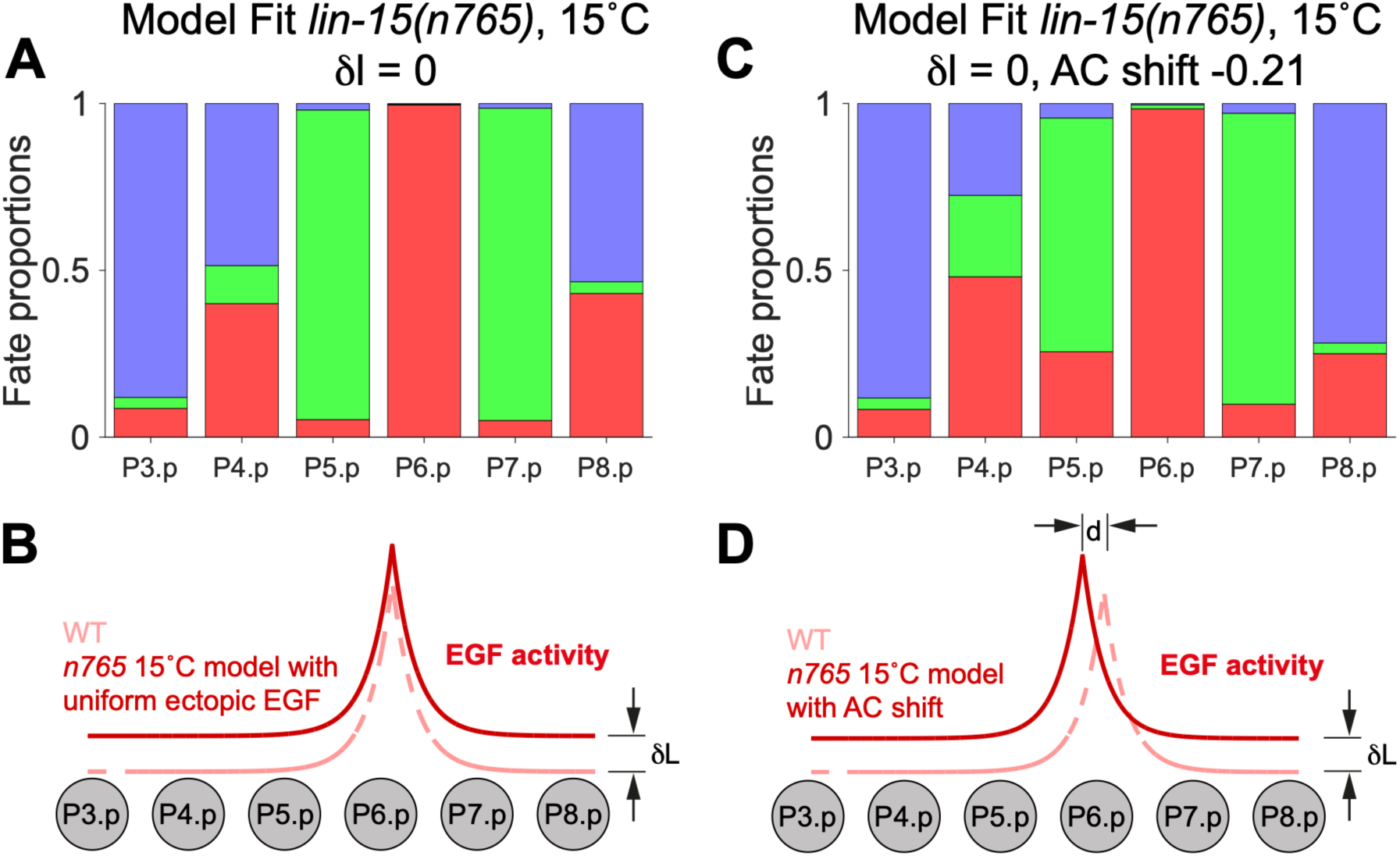
AC position shift cannot account for fate proportions in *lin-15(n765)*. (**A, B**) Fate proportions in the model fit to *lin-15(n765)* at 15°C with uniform ectopic EGF (*δl* = 0) and no AC shift (**B**), showing comparable induction levels for P4.p and P8.p (**A**). (**C**,**D**) With the AC shifted by 20% of the distance from P6.p to P5.p (**D**), the model can account for the asymmetric induction of P4.p and P8.p (**C**), but predicts only ∼ 15% induction for P3.p. compared to > 50% in experiments (compare with main Fig. 2).

**Supplementary Figure 3:**
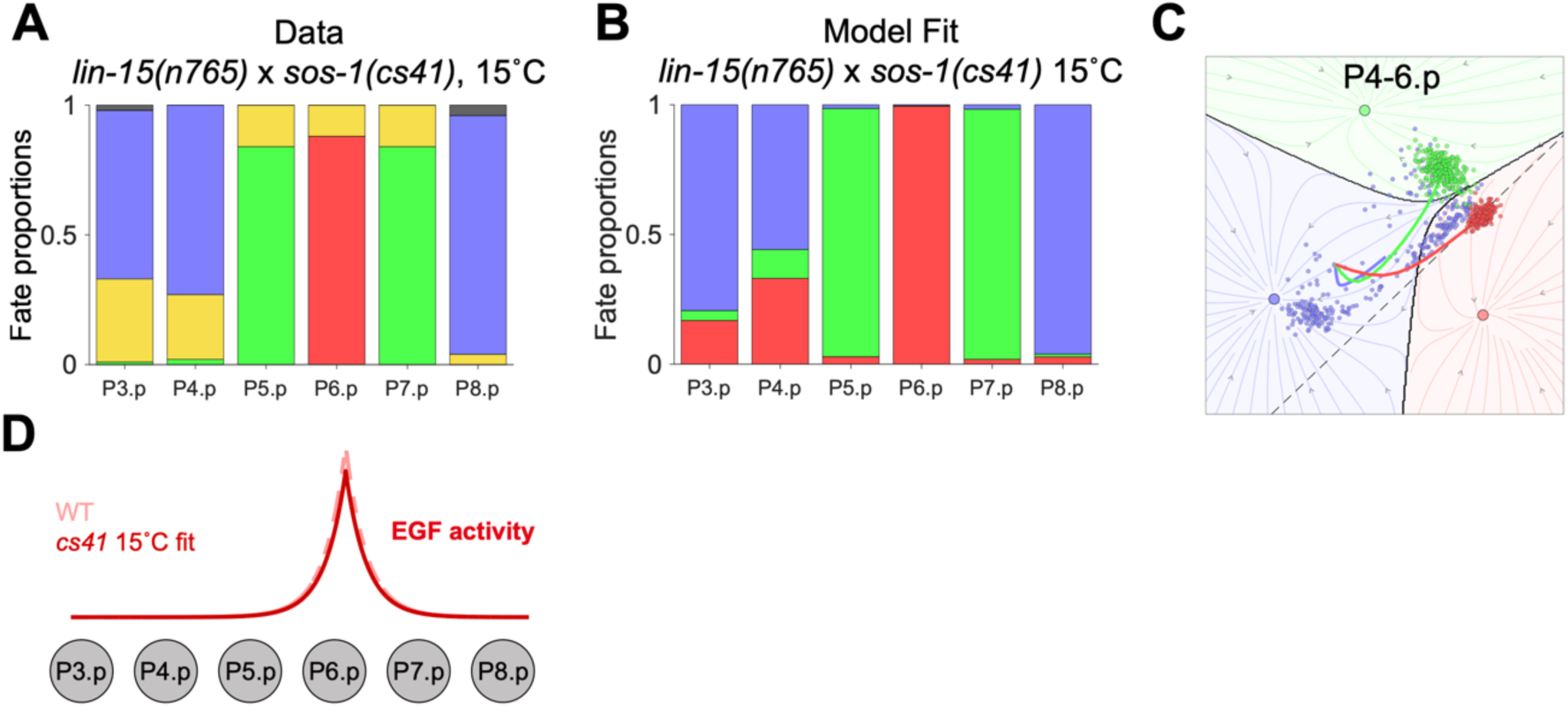
Using a cross to fit the *sos-1*(*cs41*) allele at the permissive temperature. (**A**-**B**) Experimentally observed fate proportions in the *lin-15*(*n765*) x *sos-1(cs41)* double mutant at 15°C (N=50), and the corresponding fate proportions (**B**) and fate trajectories (**C**) in the model. Ectopic EGF levels in the model are taken from our fit to *lin-15(n765)* alone (see main Fig. 2), and the fold-change in EGF activity associated with *sos-1(cs4)* (**D**) is inferred from the phenotype of the cross in **A**.

**Supplementary Figure 4:**
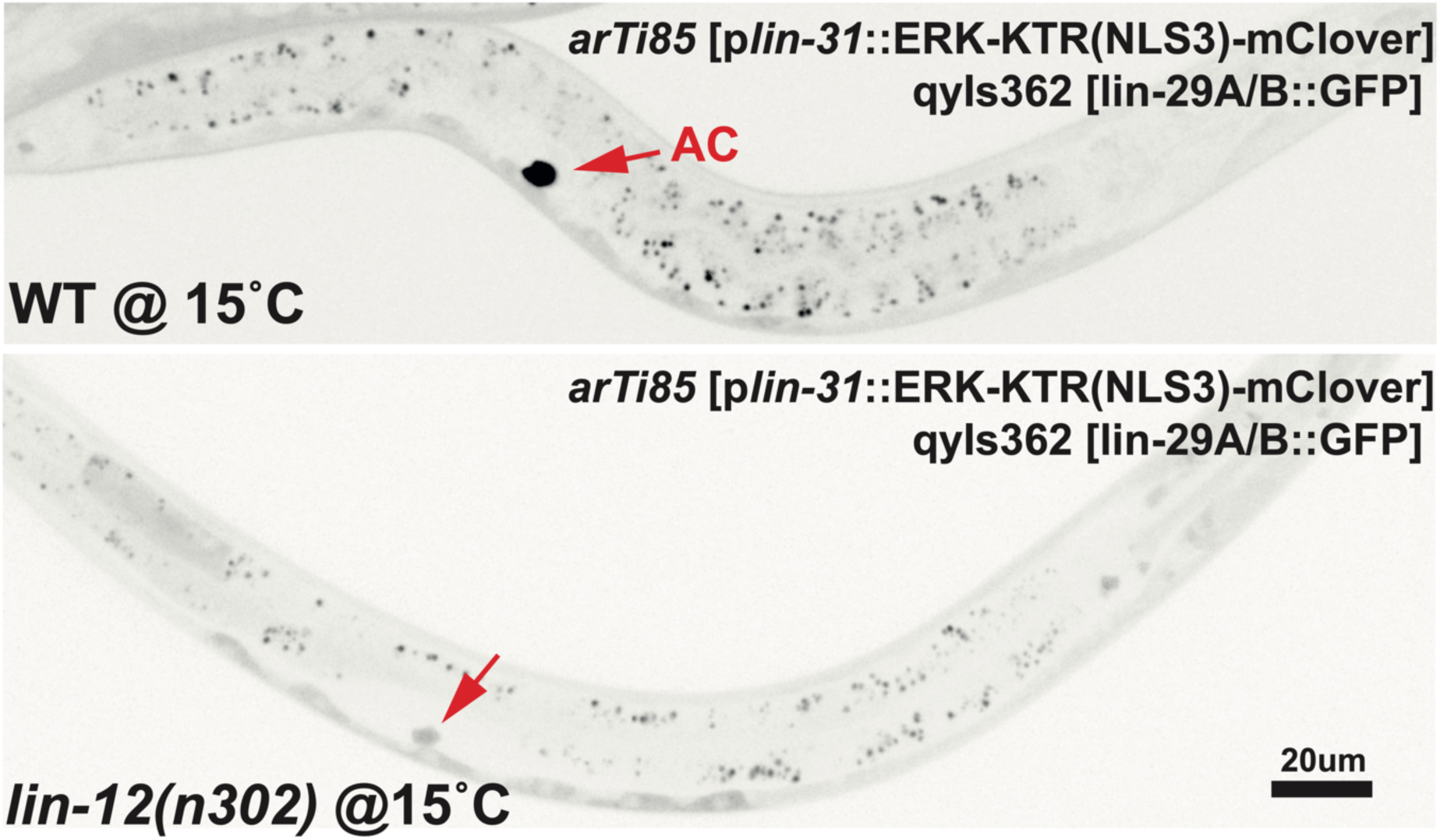
*lin-12(n302)* animals at 15°C transiently express the AC marker *qyls362* until the late L2 stage. (Top) WT late L2 stage animal with bright expression of *qyIs362* (arrow). (Bottom) *lin-12(n302)* late L2 stage animal with faint but visible expression of *qyIs362*. Both images were taken at equal exposure time and laser power.

**Supplementary Figure 5:**
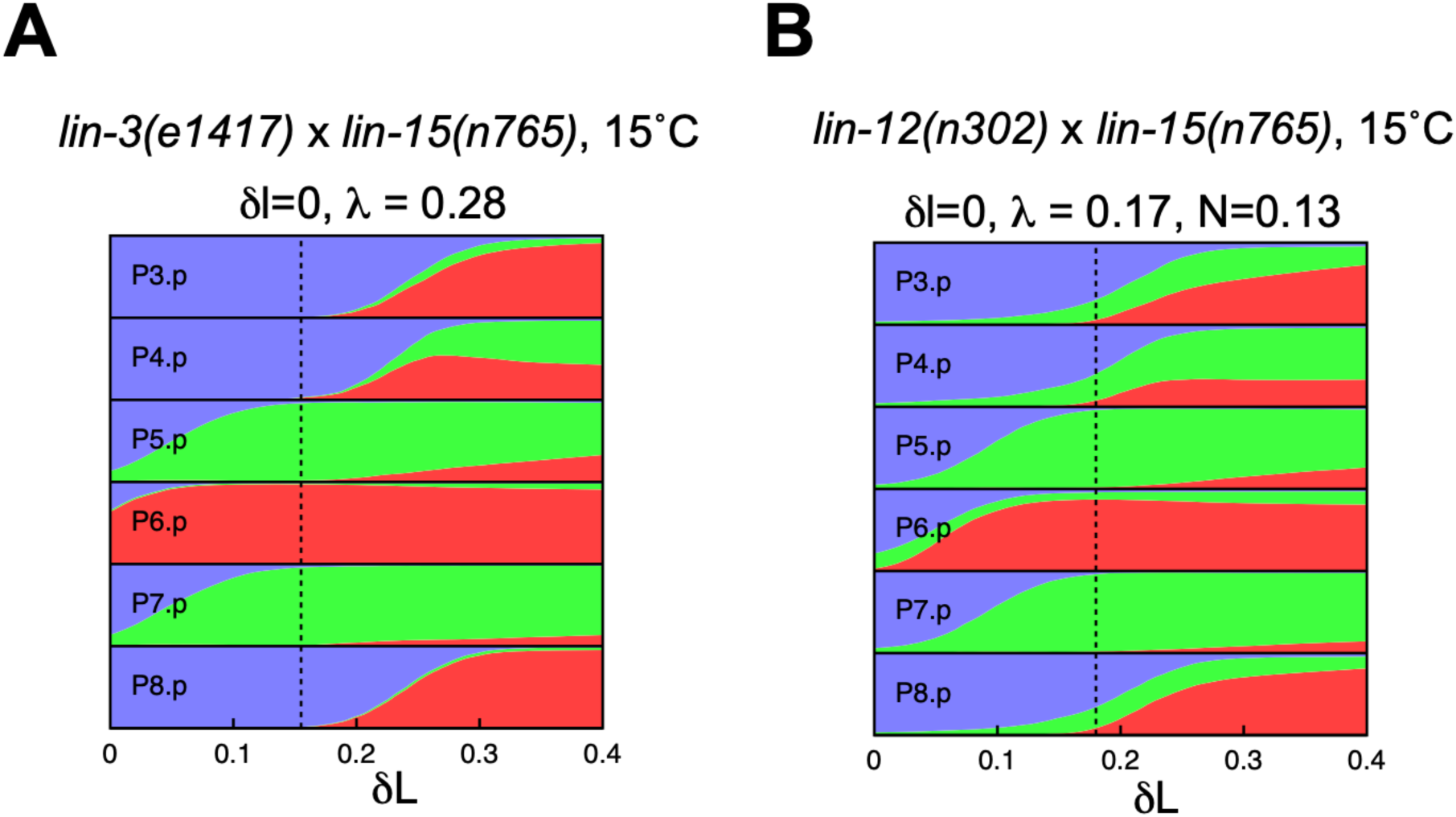
(**A,B**) Fate proportions in the model as a function of ectopic EGF level *δL* with uniform ectopic EGF (*δl* = 0), predicted for *lin-3(e1417)* x *lin-15(n765)* at 15°C (**A**) and *lin-12(n302)* x *lin-15(n765)* at 15°C. Dashed lines show our fits for the ectopic EGF level in *lin-15(n765)* at 15°C.

**Supplementary Figure 6:**
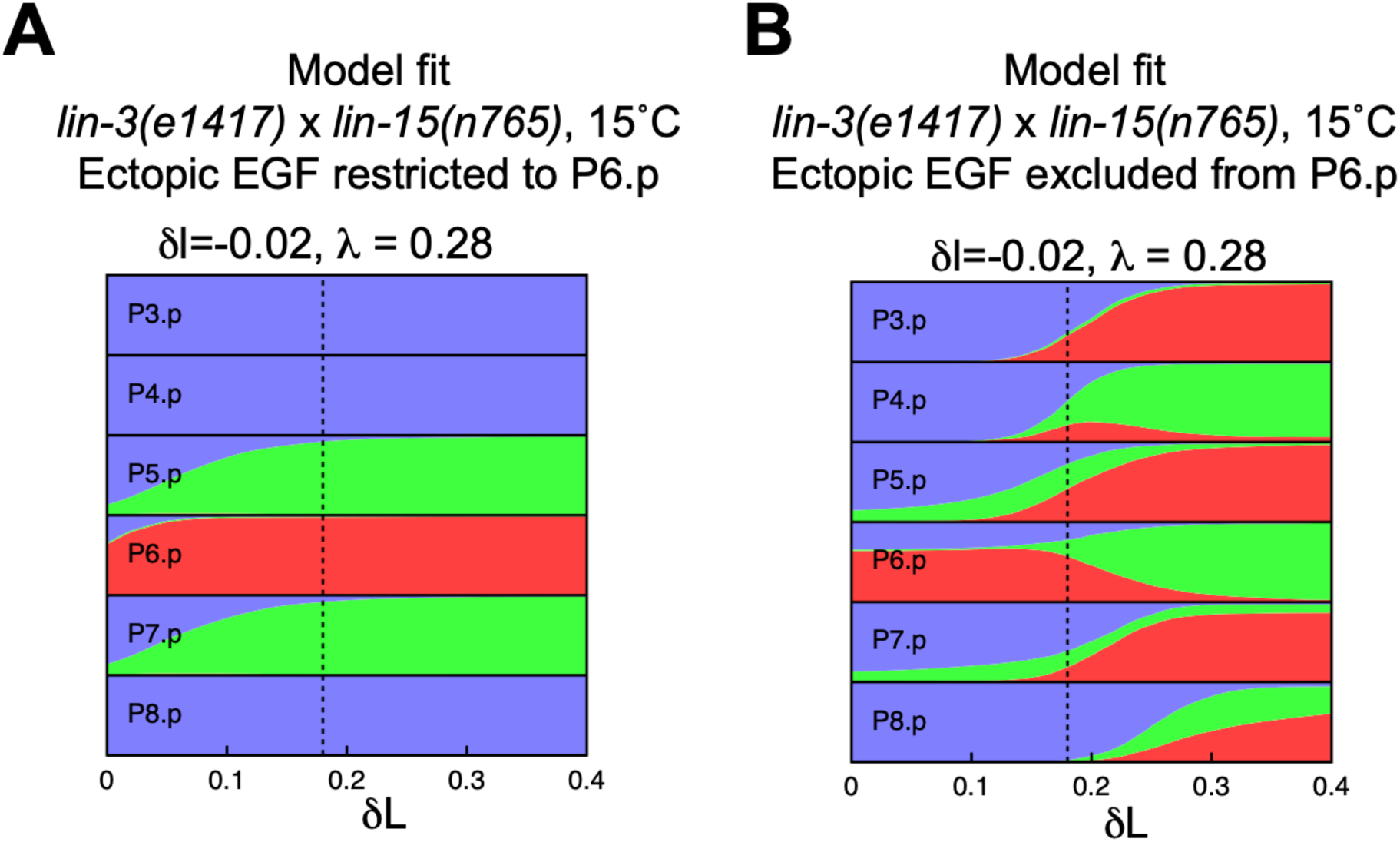
Contributions of direct vs. indirect signaling to rescue of *lin-3(e1417)* by ectopic EGF. (**A**,**B**) Predicted fate proportions in the model as a function of ectopic EGF level in the *lin-3(e1417)* background, with ectopic EGF restricted to P6.p (**A**) or excluded from P6.p (**B**), to be compared with main Fig. 4A (ectopic EGF on all cells). Dashed lines show our fit for the ectopic EGF level in the *lin-15(n765)* mutant at 15°C. In the absence of ectopic EGF signaling in P5,7.p (**A**), the stronger Notch signal received from P6.p (which itself receives a higher EGF signal) is sufficient to account for the near-complete rescue of P5-7.p to their WT fates, as seen in simulations with unrestricted ectopic EGF (main Fig. 4A-C) and in experiments (main Fig. 4D). By contrast, in the absence of additional Notch signaling from P6.p (**B**), the ectopic EGF received directly by P5,7.p themselves only partially rescues induction of P5,7.p (**B**), and results in the induction of 1° as well as 2° fates.

**Supplementary Figure 7:**
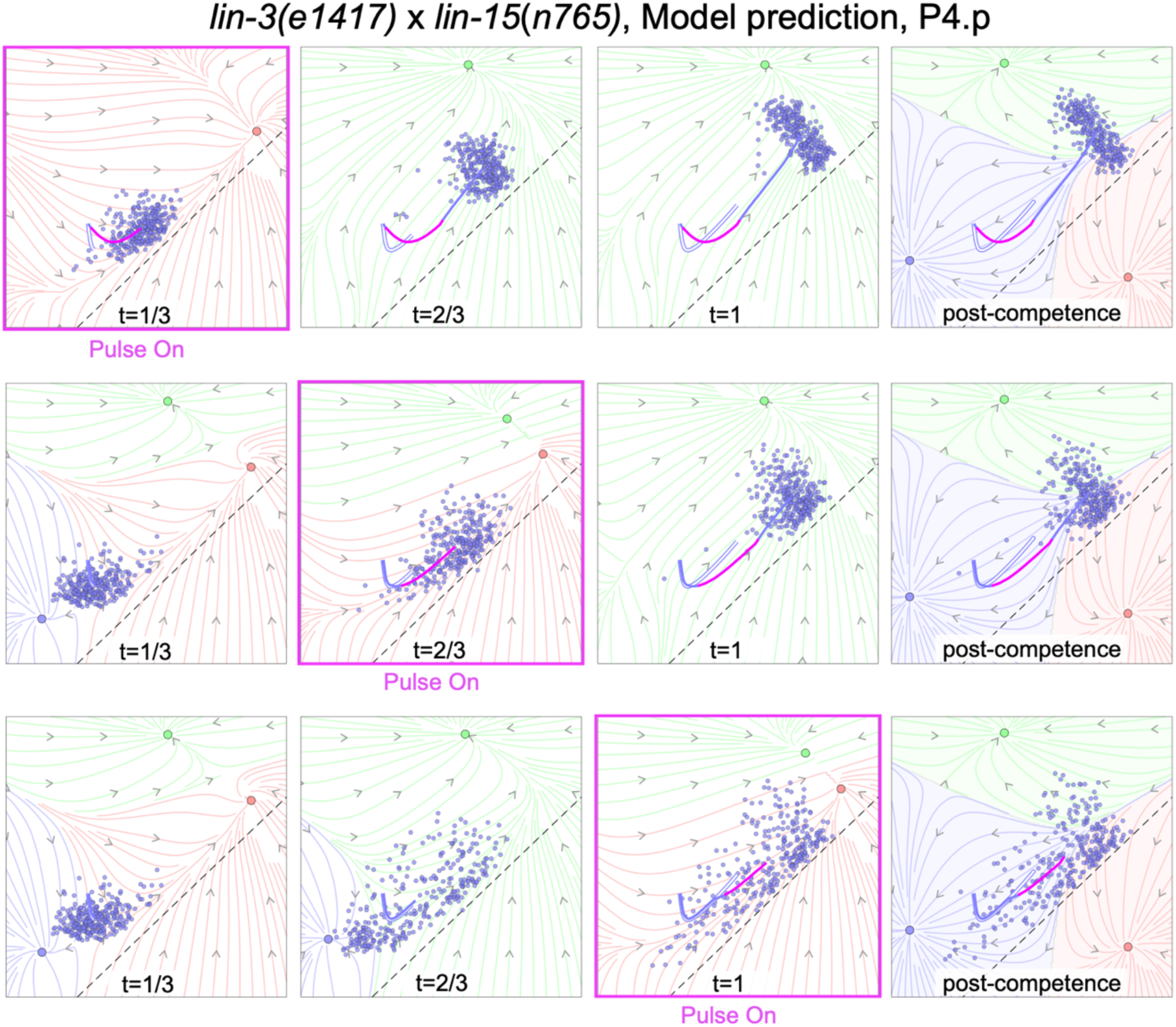
Predicted fate trajectories for P4.p in *lin-3(e1417)* x *lin-15(n765)* for early (top row), mid (middle row) and late (bottom row) temperature shifts. Flow fields determined as described in Fig. 5C, based on fits for reduced EGF from the AC in *lin-3(e1417)*, and ectopic EGF in *lin-15(n765)*. The portion of the average trajectory corresponding to the pulse is highlighted in magenta, and the average trajectory in the absence of a pulse is shown as a white curve with a blue outline.

**Supplementary Figure 8:**
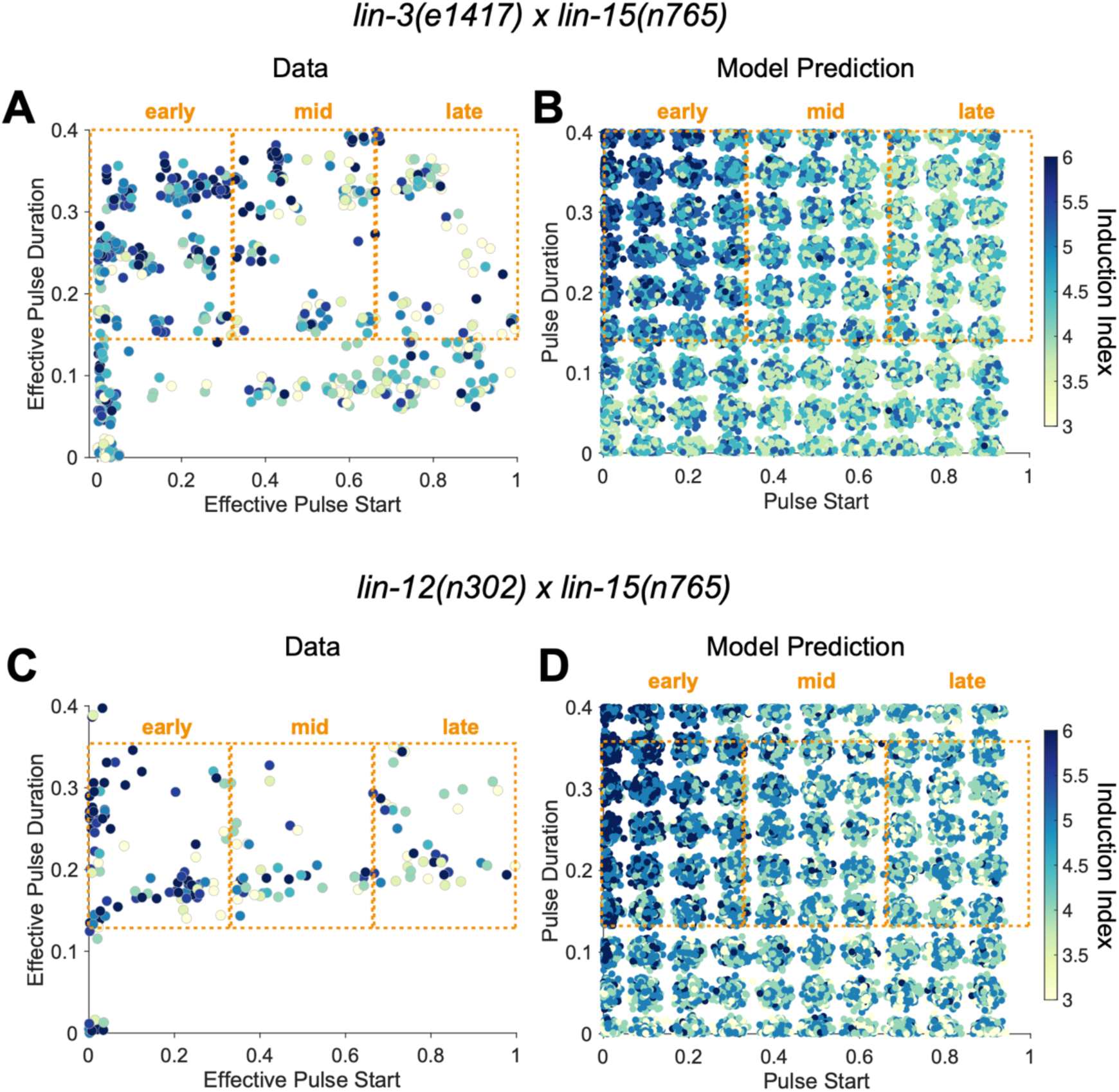
(**A,B**) Experimentally observed vulval induction levels (**A**; N=610) and model predictions (**B**) as a function of pulse start time and duration in the *lin-3*(*e1417)* x *lin-15*(*n765)* double mutant. Notice that induction levels decrease on average for pulses occurring later in the competence period. (**C**,**D**). Same as (**A**,**B**) for the *lin-12(n302)* x *lin-15(n765)* double mutant (N=309). Effective start times and durations in (**A**,**C**) are normalized to the duration of the competence period (see Materials & Methods). Dashed orange rectangles indicate which animals were included in the early-, mid-, and late-pulsed populations in main Figs. 5 and 6.

**Supplementary Figure 9:**
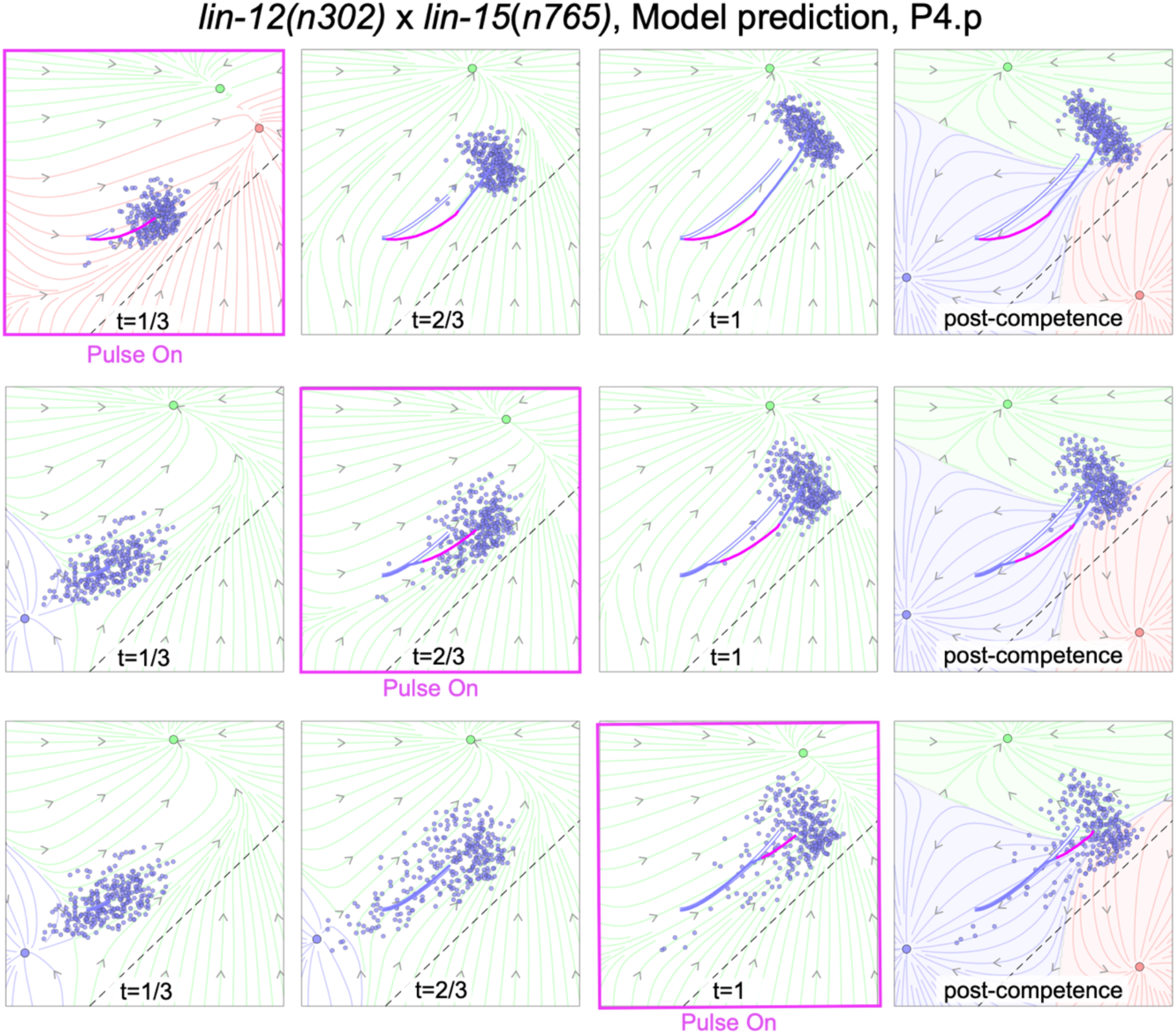
Predicted fate trajectories for P4.p in *lin-12(n302)* x *lin-15(n765)* for early (top row), mid (middle row) and late (bottom row) temperature shifts. Flow fields determined as described in Fig. 5C, but based on fits for residual EGF from the AC and constitutive Notch activity in *lin-12(n302)*, and ectopic EGF in *lin-15(n765)*. The portion of the average trajectory corresponding to the pulse is highlighted in magenta, and the average trajectory in the absence of a pulse is shown as a white curve with a blue outline.

## Supplementary Movie Legends

**Supplementary Movie 1: Dynamics of P4-6.p of an individual realization of the landscape model, during and after the competence period.** Shaded areas show the three fate valleys in the absence of signaling (see t = 0 for flow lines). Colored lines and arrows indicate instantaneous flow lines, showing how EGF and Notch signals (red and green arrows) tilt the default landscape. Once P6.p comes close the dashed line, its Notch-ligand production induces both autocrine (in P6.p) and paracrine (in P5.p) Notch signaling (growing green arrow), diverting both P5.p and P6.p upwards. For P5.p. this eventually creates a fixed point in the 2° fate area, whereas for P6.p the strong EGF signaling from the AC preserves a single fixed point in the 1° fate area. After the competence period (t > 1), we assume the default landscape which eventually distributes each cell into one of the three fate valleys

**Supplementary Movie 2: Model dynamics for WT parameters for P4-6.p** (100 realizations). The trajectories of P4-6.p are represented by blue, green, and red, respectively (according to their WT fates) and overlaid on the flow field in the absence of any signals, which applies at the end of the competence period and determines fates. The cell trajectories begin from a common point and are averaged over the noise, except at the final time.

**Supplementary Movie 3: Model dynamics for our fit to *lin-15*(*n765*) at 15°C for P3-6.p** (100 realizations). The trajectories of P3-6.p are represented by magenta, blue, green, and red, respectively.

**Supplementary Movie 4: Model dynamics for our fit to *sos-1* (*cs41*) at 15°C for P3-6.p** (100 realizations), determined from its cross to *lin-15*(*n765*). The trajectories of P3-6.p are represented by magenta, blue, green, and red, respectively. This condition is at 0.85 WT EGF signaling levels and predicted to have WT induction.

**Supplementary Movie 5: Model dynamics for our fit to *sos-1* (*cs41*) at 25°C for P3-6.p** (100 realizations). The trajectories of P3-6.p are represented by magenta, blue, green, and red, respectively.

**Supplementary Movie 6: Model dynamics for our fit to *lin-15*(*n765*) at 25°C for P3-6.p** (100 realizations). The trajectories of P3-6.p are represented by magenta, blue, green, and red, respectively.

**Supplementary Movie 7: Model dynamics for our fit to *lin-12*(*n302*) at 15°C for P3-6.p** (100 realizations). The trajectories of P3-6.p are represented by magenta, blue, green, and red, respectively.

**Supplementary Movie 8: Model dynamics for our fit to *lin-12*(*n302*) at 25°C for P3-6.p** (100 realizations). The trajectories of P3-6.p are represented by magenta, blue, green, and red, respectively.

**Supplementary Movie 9: Model dynamics for our prediction for *lin-3*(*e1417*) x *lin-15*(*n765*) at 15°C for P3-6.p** (100 realizations). The trajectories of P3-6.p are represented by magenta, blue, green, and red, respectively. Note the near complete rescue of P5.p and P6.p towards their WT fates.

**Supplementary Movie 10: Model dynamics for our prediction for *sos-1*(*cs41*) x *lin-15*(*n765*) at 15°C for P3-6.p** (100 realizations). The trajectories of P3-6.p are represented by magenta, blue, green, and red, respectively. Note the near complete rescue of P5.p and P6.p towards their WT fates.

**Supplementary Movie 11: Model dynamics for our prediction for *lin-12*(*n302*) x *lin-15*(*n765*) at 15°C for P3-6.p** (100 realizations). The trajectories of P3-6.p are represented by magenta, blue, green, and red, respectively.

**Supplementary Movie 12: Model dynamics for our prediction for *lin-12*(*n302*) x *lin-15*(*n765*) at 25°C for P3-6.p** (100 realizations). The trajectories of P3-6.p are represented by magenta, blue, green, and red, respectively.

**Supplementary Movie 13: Model dynamics for our prediction for *lin-3*(*e1417*) x *lin-15*(*n765*) P4-6.p and P8.p for early, mid, and late temperature shifts (pulses of elevated EGF signaling)** (100 realizations). The average trajectories of P4-6.p, P8.p are represented by blue, green, red, and magenta lines respectively. White lines with blue, green, red, and magenta outline indicate average trajectories of P4-6.p, P8.p if no pulse was applied (15°C constant temperature).

**Supplementary Movie 14: Model dynamics for our prediction for *lin-12*(*n302*) x *lin-15*(*n765*) P4-6.p and P8.p for early, mid, and late temperature shifts** (100 realizations). The average trajectories of P4-6.p, P8.p are represented by blue, green, red, and magenta lines respectively. White lines with blue, green, red, and magenta outline indicate average trajectories of P4-6.p, P8.p if no temperature shift was applied (15°C constant temperature).

